# Linking spatial drug heterogeneity to microbial growth dynamics in theory and experiment

**DOI:** 10.1101/2024.11.21.624783

**Authors:** Zhijian Hu, Yuzhen Wu, Tomas Freire, Erida Gjini, Kevin Wood

## Abstract

Diffusion and migration play pivotal roles in microbial communities - shaping, for example, colonization in new environments and the maintenance of spatial structures of biodiversity. While previous research has extensively studied free diffusion, such as range expansion, there remains a gap in understanding the effects of biologically or physically deleterious confined environments. In this study, we examine the interplay between migration and spatial drug heterogeneity within an experimental meta-community of *E. faecalis*, a Gram-positive opportunistic pathogen. When the community is confined to spatially-extended habitats (‘islands’) bordered by deleterious conditions, we find that the population level response depends on the trade-off between the growth rate within the island and the rate of transfer into regions with harsher conditions, a phenomenon we explore by modulating antibiotic concentration within the island. In heterogeneous islands, composed of spatially patterned patches that support varying levels of growth, the population’s fate depends critically on the specific spatial arrangement of these patches - the same spatially averaged growth rate leads to diverging responses. These results are qualitatively captured by simple simulations, and analytical expressions which we derive using first-order perturbation approximations to reaction-diffusion models with explicit spatial dependence. Among all possible spatial arrangements, our theoretical and experimental findings reveal that the arrangement with the highest growth rates at the center most effectively mitigates population decline, while the arrangement with the lowest growth rates at the center is the least effective. Extending this approach to more complex experimental communities with varied spatial structures, such as a ring-structured community, further validates the impact of spatial drug arrangement. Our findings suggest new approaches to interpreting diverging clinical outcomes when applying identical drug doses and inform the possible optimization of spatially-explicit dosing strategies.

**Author summary:** In this study, we develop an automated platform to experimentally investigate short-term population growth and migration dynamics under spatial drug heterogeneity. Our findings reveal that the collective spatial response of the population can vary significantly, even with the same migration rate and averaged drug dose, due to different spatial drug arrangements. By constructing a simple reaction-diffusion model, we observed that simulated short-term spatial growth rate closely matches the experimental data. Furthermore, this short-term spatial growth rate aligns well with the long-term spatial growth rate, defined by the largest eigenvalue, as the spatial system quickly enters the equilibrium growth state. Using concepts from perturbation theory, we derived an analytical relationship between the boundary diffusion effect, homogeneous growth effect, and heterogeneous effect. Our results highlight that in spatially-extended habitats, the spatial growth response is an emergent property. The bacterial population quickly enters equilibrium growth, suggesting that the spatial growth rate measured at an ecological scale may be used to predict resistance evolutionary behavior.

## Introduction

The bacteria infection and resistance has become a worldwide health problem [1–4]. Starting from the 1940s, the use of antibiotics has been one of the most powerful tools in taming microbial pathogens [5]. In laboratory studies, people usually study the response of well-mixed population to the drug [6–16, 16–22]. However, the living environments in the human body are usually spatially-relevant and heterogeneous; evidence has been found in the gut and tumors [23, 24]. Some previous studies show that, spatial gradients in drug concentration dramatically accelerated resistance evolution [12, 25–41]; under drug gradients the slow bacteria can be more resistant and thus give us a trade off [42]. On a larger spatial scale, like organ level, spatial drug heterogeneity is also found between lung and gut, which elevates the bacteria survival and resistance [43]. Some theories have also been developed to find out the non-monotonic evolution behavior under spatial drug heterogeneities [44] and genotypic fitness landscapes [45]. Diverging clinical outcomes—whether the pathogen population is cleared or not—arise due to the complex dynamics of bacterial populations [46]. How spatial drug heterogeneity generally shapes bacterial growth dynamics and the spatial collective response of populations, thus altering treatment outcomes, is still not fully understood.

Most experimental studies have focused on monotonic drug gradients [25, 42, 47, 48, 48, 49], 2-well drug sanctuary environments [50–52], and the range expansion of surface-attached biofilms [53–55]. However, the drug environment within the human body, like in the gut, is typically more non-monotonically heterogeneous and fluctuating [56]. In addition to forming condensed biofilms, bacteria often exist in a planktonic form, living in a liquid environment. While many studies focus on range expansion both theoretically and experimentally [57, 58], they frequently assume a free-diffusion model that requires infinite free space—an assumption that is unrealistic in natural or human environments where physical or biological confinement is common. In human bodies, bacteria or tumor cells are often confined by tissues, vessels, or immune and acidic environments, like scattered islands surrounded by the sea in island geography. These confined boundaries can be deleterious - migrating out of the confined regions can be deadly. For example, cancer patients who experience radiation therapy have radiation regions where bacteria will die. These deleterious regions are also common when the regions are surgically removed, or nutrition severed. Greater attention needs to be given to confined environments and bacteria migration between these deleterious boundaries. Furthermore, there is a lack of investigation into ecological time scale dynamics under spatial drug heterogeneity. Clinically, treatment-induced resistance [59] is often a result of drug heterogeneity and inefficient bacterial clearance, occurring at an ecological level before the onset of evolution. Therefore, despite its simplicity, studying short-term bacterial responsive dynamics under controllable non-monotonic spatial heterogeneity and a deleterious confined environment, may be key to better understanding diverging clinical outcomes and pathogen recurrence in hospitals and patients.

In this study, we utilize the wild-type Gram-positive opportunistic pathogen *Enterococcus faecalis* as our experimental bacterium. *E. faecalis* thrives in the human gastrointestinal tract and is responsible for numerous clinical infections, including 5 to 15 percent of cases of infective endocarditis and urinary tract infections [60–64]. To investigate how spatial drug heterogeneity affects bacterial population dynamics under a deleterious confined environment, we employ a specialized, island-like experimental system with 2 absorbing boundaries, facilitated by a pipetting robot. First, we demonstrate that bacterial migration in a confined environment yields distinct population survival outcomes when the drug is homogeneously distributed, indicating that system size and drug concentration are critical factors, presenting a trade-off relationship. In environments with arbitrary non-monotonic drug gradients, our findings reveal that spatial drug heterogeneity significantly alters population dynamics, with the effects of different spatial arrangements being as substantial as, or even greater than, the drug effect itself. Furthermore, we observe that increasing the drug amount and migration rate leads to markedly different outcomes for different selected spatial arrangements. We hypothesize that spatial drug arrangements, combined with boundary effects, create varying levels of spatially favorable habitats. These results are qualitatively captured by simple simulations and analytical expressions derived using first-order perturbation approximations to reaction-diffusion models with explicit spatial dependence. Among all possible spatial arrangements, our theoretical and experimental findings reveal that central drug-free habitats most effectively mitigate population decline, while central drug habitats are the least effective. This aligns with the previous theoretical finding on optimal fragmentation of invasion in heterogeneous habitats [65]. Extending this approach to more complex experimental communities with varied spatial structures, such as a ring-structured community, further validates the impact of spatial drug arrangement. Our findings build a direct link between theoretical predictions and experimental validations of bacterial population response under spatial drug heterogeneity. It may provide new approaches to interpreting diverging clinical outcomes when applying identical drug doses and inform possible optimizations of personalized dosing strategies.

### Experimental set-up for bacterial growth and diffusion dynamics under drugs

To study the effect of short-term diffusion, we let the E. faecalis bacterial population migrate to its nearest neighbor along the 1D space, by each row of the 96 well plates (Figure 1B). The system size is determined by the number of wells selected from a total of 12 wells per row. For each well plate, we can then have 8 replicates for the data analysis. Migration is achieved by exchanging small volumes of growth media between adjacent wells. The experiments were started with a uniform initial population density profile, and after each diffusion cycle the cell density was measured by the plate reader. To ensure spatial drug homogeneity, we administer a uniform drug concentration *D* to each well. In contrast, spatial drug heterogeneity is achieved by varying drug concentrations across wells. For simplicity and without loss of generality, we utilize a combination of drug-free wells and high-drug wells beyond minimal inhibitory concentration(MIC) that completely inhibit bacterial growth. Different spatial drug arrangements are then created by permuting these drug-free and high-drug wells. Linezolid(LZD) are used in this study. Drugs are preloaded in the form of media.

**Fig 1.**
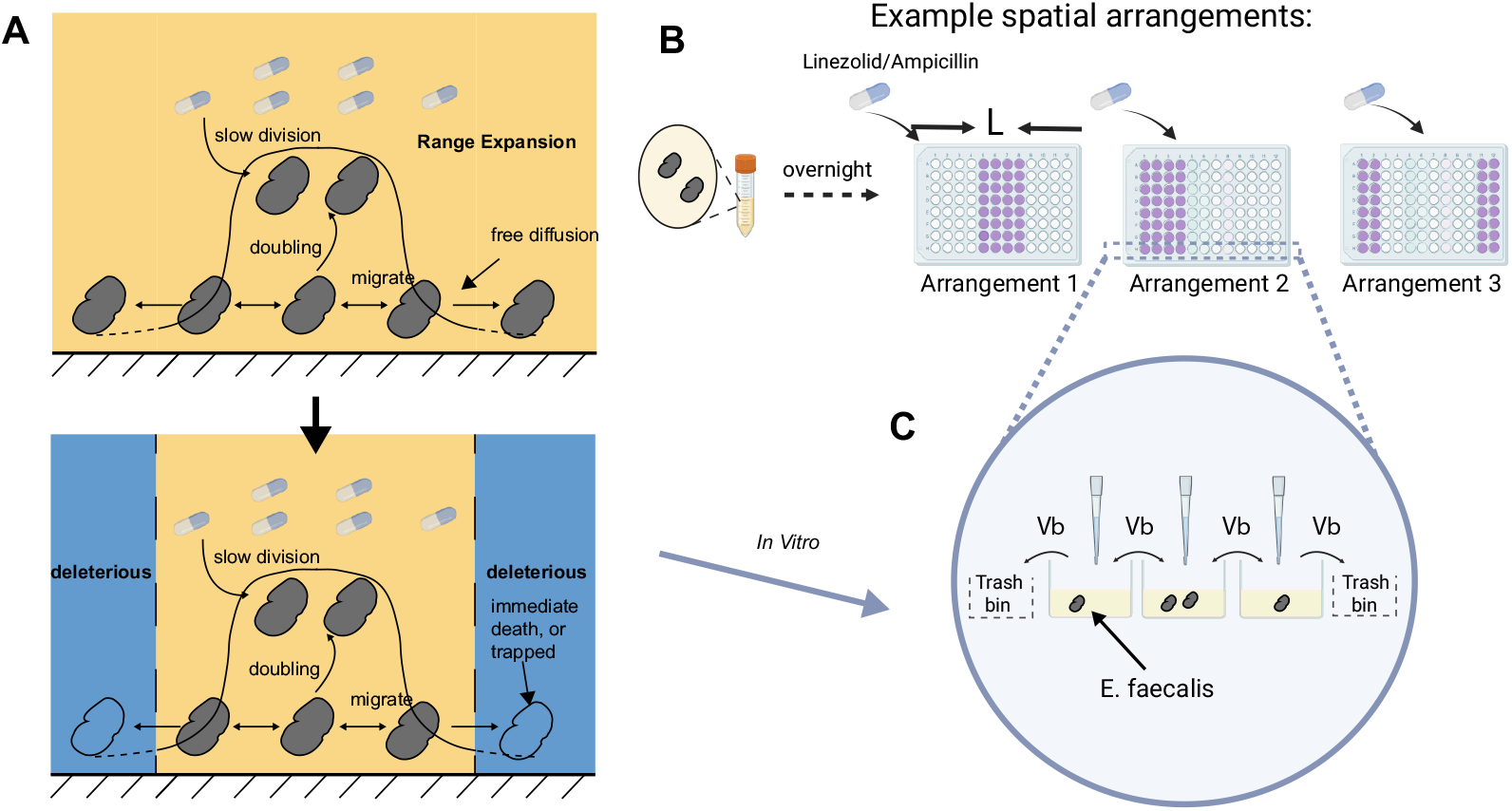
Schematic of growth-migration dynamics in a deleteriously confined environment, and experimental design. **A**. The illustration above shows a bacterial population growing and migrating freely in an unbounded environment with spatial drug heterogeneity. The following illustration represents the population proliferating and migrating in a deleteriously confined environment. **B, C** The *E. faecalis* was grown overnight and then diluted 1:1 into different 96-well plates with a spatially uniform density profile, but different drug concentrations *D* and spatial arrangements, at initial time *T* = 0. After every growth cycle of *T* h, bacteria migration was performed by transferring the same amounts *V b* of bacteria liquid to both neighboring wells along the columns by the OT-2 pipetting robot; *V* is the total volume per well and usually is 200*µL*; b is the transferred fraction. Bacteria at 2 boundary wells were taken out at the same volume *V b*. Cycles would be repeated after the growing density curve had equilibrated. Usually it’s ∼ 8 times. Cell density profiles were measured by plate reader exactly before the next migration/volume transfer.

To maintain a deleterious environment, for the bacteriostatic drug Linezolid that only inhibits bacteria growth, at the end of each diffusion cycle, *b* fraction of the total volume *V* will be taken out from the 2 boundary wells, and then they will be re-supplied with drug-free media or high-drug media, depending on the spatial arrangement of the drug, to compensate for the media and drug loss. This also helps us keep the drug distribution roughly unchanged. Therefore, we can ignore the drug diffusion dynamics.

To ensure that the bacterial population remains within the exponential growth phase, we control our experiment to be of limited duration, focusing on an ecological time scale. Specifically, the total experiment time is set to span 8 cycles (each cycle being either 2 or 4 hours). This approach helps to maintain the integrity of the spatial drug response and minimizes the complexity introduced by extended experiments. In particular: 1. The short duration prevents the emergence of mutations and resistance evolution in the bacterial population. 2. The population is away from carrying capacity. 3. Drug diffusion effects are kept minimal, preserving the initial drug concentration distribution. This experimental setup allows us to isolate and observe the targeted spatial and collective drug responses without the interference of longer-term evolutionary and diffusion dynamics.

## Mathematical model formulation

At the continuous limit, this experimental system is actually a simplified Fisher-KPP equation with 2 absorbing boundaries

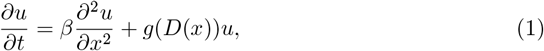

with *u*(0, *t*) = 0, *u*(*L, t*) = 0 to describe the deleterious environment outside of our spatially-extended habitats. *u*(*x, t*) is the cell density at position(well) *x* at time *t. g*(*D*(*x*)) the growth rate at position *x* corresponding to the drug concentration *D. β* is the diffusion or migration rate and *L* is the length of the wells used in a well plate. In a discrete version of this reaction-diffusion equation, *L* represents the number of total wells used, as the spatially-extended habitats. By comparing it with the discrete dynamical equation of the experimental protocol, we can connect *g, β* with our experimental parameters,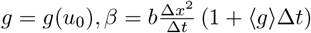, where Δ*x* = 1 well, Δ*t* = 0.25/0.5h (Supplementary Information).

The system will experience a transient, fluctuating population change over space at initial times. After the system is equilibrated and entering a stable growth or decline phase, the survival criterion is given by the largest eigenvalue of the operator (Supplementary Information)

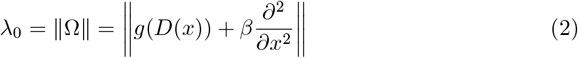

For our model, the largest eigenvalue can be separated into 2 terms which describe the growth (*g*_*eff*_) and boundary diffusion effect 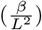. The bacteria gives a declining response when

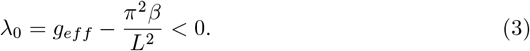

For the homogeneous case, *g*_*eff*_ = ⟨*g*⟩ = *g*(*D*) is exactly the growth rate corresponding to the drug concentration; for the heterogeneous case, *g*_*eff*_ can be approximated by the 1st-order perturbation theory, as *g*_*eff*_ = ⟨*g*⟩ + ⟨*u*_0_|*δg*|*u*_0_⟩ (Supplementary Information), where 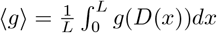 describes the spatial homogeneous effect, and 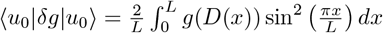 describes the spatial heterogeneous effect. 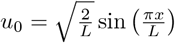 is the eigenvector corresponding to the larges unperturbed eigenvalue, and *δg* = *g*(*D*(*x*)) − ⟨*g*⟩ is the growth rate deviation. Although here we only consider the single-drug response of the homogeneous bacterial population, in our recent work, the derivation results above can be generalized to multi-strain systems under multi-drug selections with tunable spatial gradients, determining the most dominant resistant strain [66].

## Results

### Bacteria shift from growth to decline by increasing drug concentrations and boundary diffusion

A natural question to ask first, is how diffusing outside the deleterious confined environment, the boundary diffusion effect 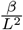, shapes the population dynamics. For simplicity, we start with homogeneous growth rates with all wells containing the same drug concentration *D* over the space, modulated by a bacteriostatic drug Linezolid. It inhibits bacterial growth but does not cause a decline in the population itself. By varying drug concentrations *D* over patches, we find that, under low drug concentrations, bacteria can adapt and thrive despite the deleterious environment, leading to an increase in cell density. This is reasonable because the uniform growth rate, which drives the increase in population density, outcompetes the boundary diffusion effect 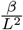, which diminishes the population. As drug concentrations *D* go higher, the bacterial growth diminishes significantly, impairing the population’s ability to reproduce sufficiently to counteract cell loss by the boundary diffusion effect. This imbalance causes the population to decline, ultimately resulting in extinction as cell density trends towards zero over a long time limit. Figure 2A depicts bacterial growth dynamics in drug-free conditions (D=0 µg/ml) and under high drug concentrations (D=8 µg/ml). As we can see, by increasing the drug concentration(Figure 2B, left panel), the spatial collective response of population shifts from growth to decline. The criterion for bacterial decline, with experimental data, is determined by comparing the final optical density(OD) to the initial ODs, as described in detail in the Supplementary Information.

**Fig 2.**
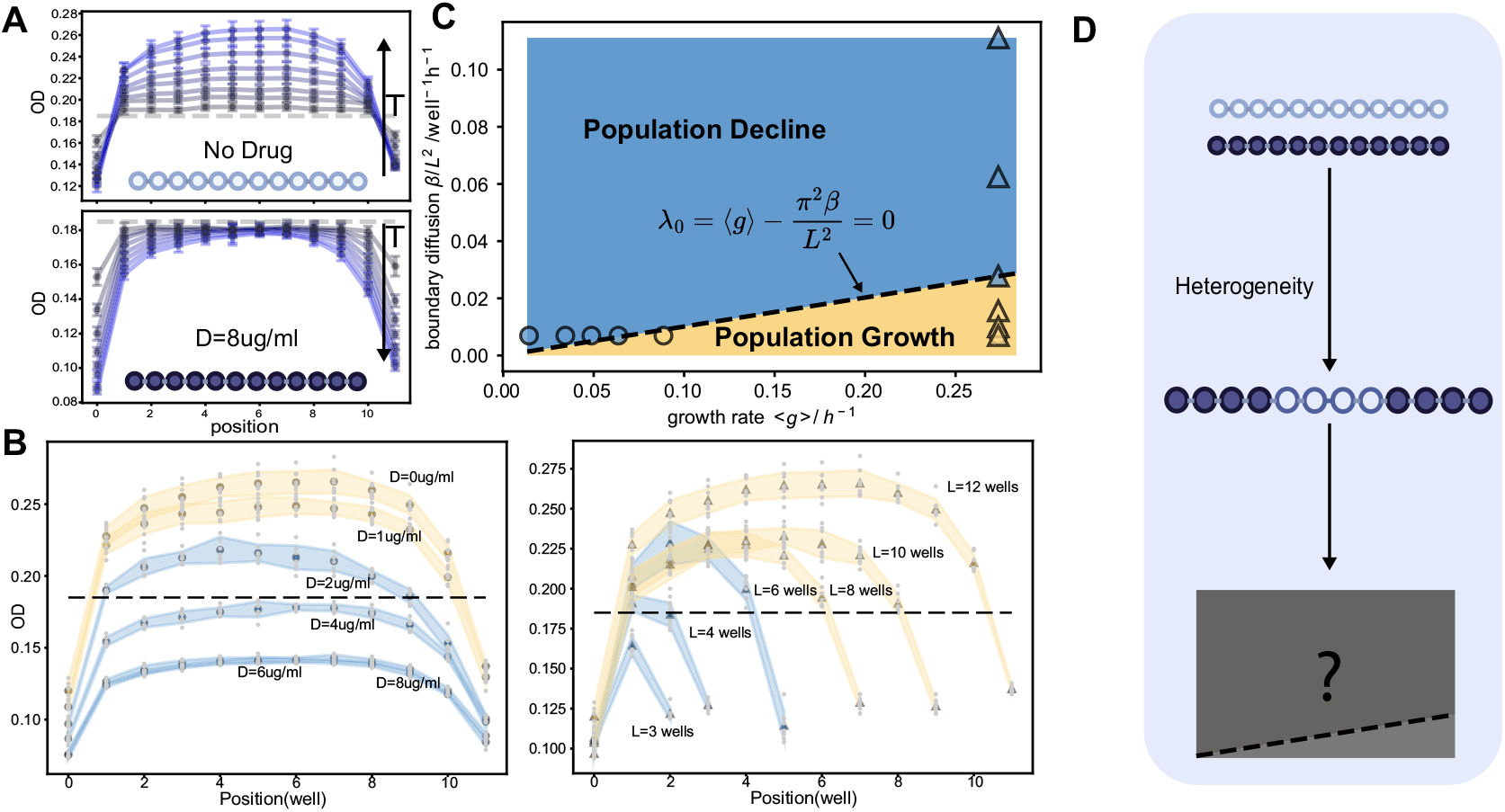
Bacteria population response (growth/decline) in different drug concentrations and migration regimes of a bacteriostatic antibiotic, Linezolid, for 8 cycles. **A**. Position-specific bacteria growing process under spatial drug homogeneity in drug-free(D=0ug/ml) regimes and high-drug(D=8ug/ml) regimes. At the bottom of each figure, a 12-well illustration with light or dark colors represents drug-free wells or drug wells. The dashed line in each figure is the initial spatial cell density at *T* = 0. Grey blue dots and curves are early cycles while light blues are late cycles. For drug-free regimes, as time increases, the curve is gradually shifting up while the spatial density curve is decreasing down for high-drug regimes. Each single curve with error bars including initial cell densities are averaged over replicates of 8 rows in the 96-well plate. **B**. Endpoint OD curves, for 6 different drug concentrations *D* (left panel, circles) and 6 system sizes *L* (right panel, triangles). The blue scattered dots represent conditions where the final bacterial population increases, while the orange dots indicate where the population decreases. The shaded regions denote error margins. **C**. A phase diagram showing the relationship between the boundary diffusion effect,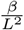, and the spatially averaged growth rate, ⟨*g*⟩.

Next we fix the homogeneous growth rate(with no drug) to investigate the boundary diffusion effect 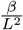, by tuning the system size *L*. Again, the population shifts from growth to decline when the system size *L* is reduced from 12 wells to 3 wells (Figure 2B, right panel), as predicted by the largest eigenvalue criterion. Thus population declines or not hinges on the trade-off between homogeneous growth rate *g*(*D*), and boundary diffusion effect 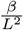. Our experimental data matches well with the growth and decline phases in the phase diagram (Figure 2C), where the transition boundary is determined by 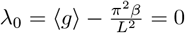. Since growth rate is homogeneous here we use ⟨*g*⟩ to replace *g*(*D*), for comparison with spatially heterogeneous growth rates. This quantitative trade-off relationship matches with the classic critical patch size result 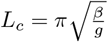, in the study of reaction-diffusion models, particularly in ecological and biological contexts [67], and may be helpful to explain the colonization of the gut microbiome in the human body [23].

### Different spatial drug arrangements modulate growth dynamics

After understanding how a deleteriously bounded environment incurs population decline, we next investigate the effect of spatial drug heterogeneities. In homogeneous drug environments, bacterial communities either grow or decline, determined by boundary diffusion and a fixed spatially averaged growth rate. However, in a spatially heterogeneous drug environment, different spatial drug arrangements may result in varying temporal growth dynamics, leading to different population outcomes, even with the same spatially averaged growth rate ⟨*g*⟩. The next question to explore is how spatial drug heterogeneity alters growth dynamics experimentally, and whether any new emerging patterns can be predicted with our simplified reaction-diffusion model (Figure 2D).

For a specific total drug amount or a fixed spatially averaged drug concentration, or growth rate ⟨*g*⟩, numerous spatial drug arrangements can be designed. For simplicity, we use *D* = 0 µg/ml and *D* = 8 µg/ml with different spatial arrangements to create a binary-heterogeneous environment, consisting of drug and non-drug wells. The spatially averaged growth rate can be represented by the number of drug wells, *n*_*D*_, while keeping the number of drug wells fixed and permuting their order for comparison.

To start, we designed 6 different drug well arrangements (See Figure 3A, I-VI): center drug-free wells (I), left-side drug-free wells (II), left edge drug-free wells (III), center drug wells (IV), left-side drug wells (V), and left edge drug wells (VI). Center drug-free wells are referred to as CH, as they have the high growth rates at the center; similarly, CL is used as a short form for center low growth rates, or center drug wells. Configurations I-III share the same number of drug wells *n*_*D*_ = 8, while configurations IV-VI share the same number of drug wells *n*_*D*_ = 4. For each group, we aim to understand how growth dynamics are influenced by different spatial arrangements and to compare the differences between groups. Figure 3B presents the temporal dynamics of these 6 examples, illustrating reshaped density curves as expected due to the spatial drug arrangements. Interestingly, a pattern emerges within these two groups: as drug-free wells are positioned closer to the center of the spatially extended habitats, the final ODs are higher (I *>* II *>* III and IV *<* V *<* VI, see Figure 3C, 3D). Populations in II and III decline, while populations in I, IV, V, and VI grow. This provides direct evidence of the spatial arrangement effect. Since there are both population growth and declines even with the same number of drug wells, these diverging outcomes don’t belong to either the growth phase or decline phase in Figure 2C, thus can’t be simply determined by the boundary 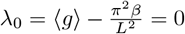. It indicates that in the 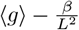 phase diagram, a new induced mixed response phase may exist, tuned by spatial weighting of the arrangements.

**Fig 3.**
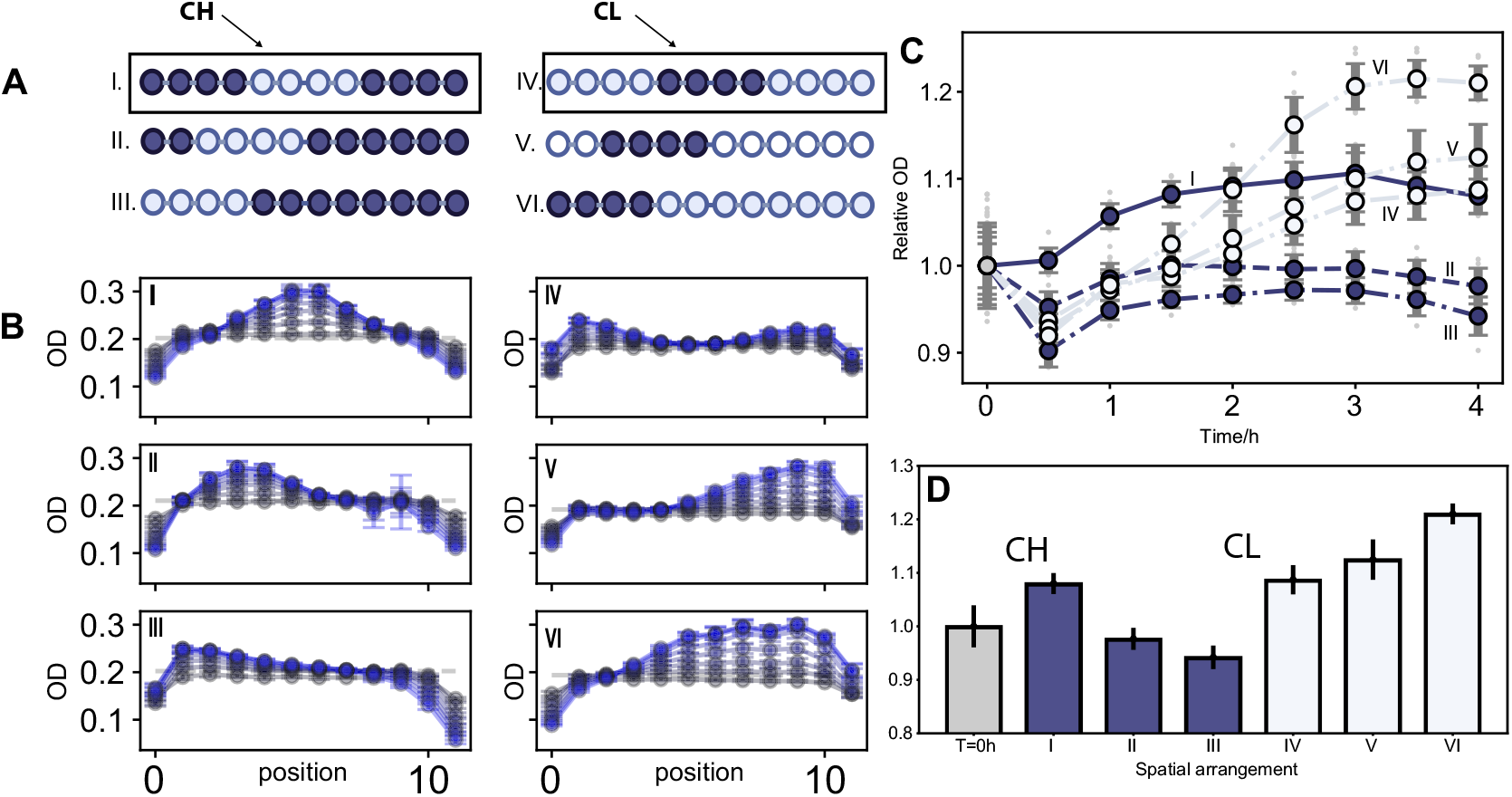
Different spatial arrangements lead to different temporal dynamics and collective responses. **A**. Six different spatial arrangements are depicted. Each column has the same number of wells with drugs, resulting in the same spatially averaged growth rate. Dark color indicates drug wells and light color indicates drug-free wells. The first column contains 8 wells with drugs; the central region initially has 4 drug-free wells with high growth rates, named Central High (CH). The second column contains 4 wells with drugs, and the central region initially has 4 high-drug wells with low growth rates, named Central Low (CL). In both CH and CL configurations, the central region is shifted two wells to the left in each subsequent arrangement (II,III) and (V,VI). **B**. The temporal dynamics of the six different spatial arrangements, with error bars corresponding to ±1 standard deviation of 8 technical replicates. Dashed line represents the initial OD at *T* = 0. Grey blue dots and curves are early cycles while light blues are late cycles. **C** and **D** present a comparison of the averaged temporal dynamics and final outcomes across six different spatial arrangements. In both panels, the y-axes represent the relative final ODs, normalized by dividing by the initial OD for comparison purposes. Dark blue represents I,II,III, while light blue represents IV,V,VI.

Although I-III, with a lower averaged growth rate, would intuitively have lower final ODs compared to IV-VI with double the amount of drug wells, our results show that the spatial arrangement with center drug-free wells (I) yields results very close to those of the spatial arrangement with center drug wells (IV) or edge drug-free wells (III). In Supplementary Information, another repeated experiment demonstrates that the population in I grows while the population in IV declines, indicating an even larger spatial arrangement advantage of CH over CL. This discrepancy of 2 different experiments may be due to fluctuations in drug concentration and temperature from day to day. Both of them can serve as supporting evidence for the hypothesis that, arrangement with the highest growth rates at the center most effectively mitigates population decline, while the arrangement with the lowest growth rates at the center is the least effective. Our simulation results (see Supplementary Information) closely match the observed temporal dynamics and population responses. These results together suggest that center drug-free wells(CH) and center drug wells(CL) may serve as the upper and lower bounds for the possible mixed phase in the 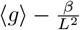 phase diagram, warranting further investigation of our model system to gather more evidence.

### Theory validates the indicated mixed phase, and explains spatial effect

For a fixed value of spatially averaged growth rate ⟨*g*⟩, we can design different strategies to assign growth rates in wells between 0 and drug-free growth rate *g*_0_, constrained by ⟨*g*⟩. To begin with, six spatial arrangement strategies were designed for comparison: Homo, OddEven, Randomized, Left, CH, and CL. As indicated by our spatial arrangement experiments, each fixed arrangement strategy induces a specific phase diagram of ⟨*g*⟩ and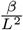 (see Figure 4A). The “Homo” strategy is always distributing the spatially averaged growth rate ⟨*g*⟩ to each well, showing a consistent phase diagram with Figure 2C, while the “OddEven” strategy, which distributes growth rates in odd-numbered wells first until they are all saturated at *g*_0_ and then in even-numbered wells, produces a more curved boundary between population growth and decline. The “Randomized” strategy randomly assigns growth rate values to each well, and yields a phase diagram nearly similar to that of homogeneity. The “Left” strategy, which assigns growth rates from left wells to right wells sequentially, results in a distinct pattern. “CH” strategy is distributing growth rates at center wells first, and “CL” strategy is distributing growth rates at edge wells first. Comparing all six strategies, CH results in the largest phase of population growth, while CL results in the largest region of population decline, and they are bounding all 6 different spatial arrangement strategies (see Figure 4A and 4B).

**Fig 4.**
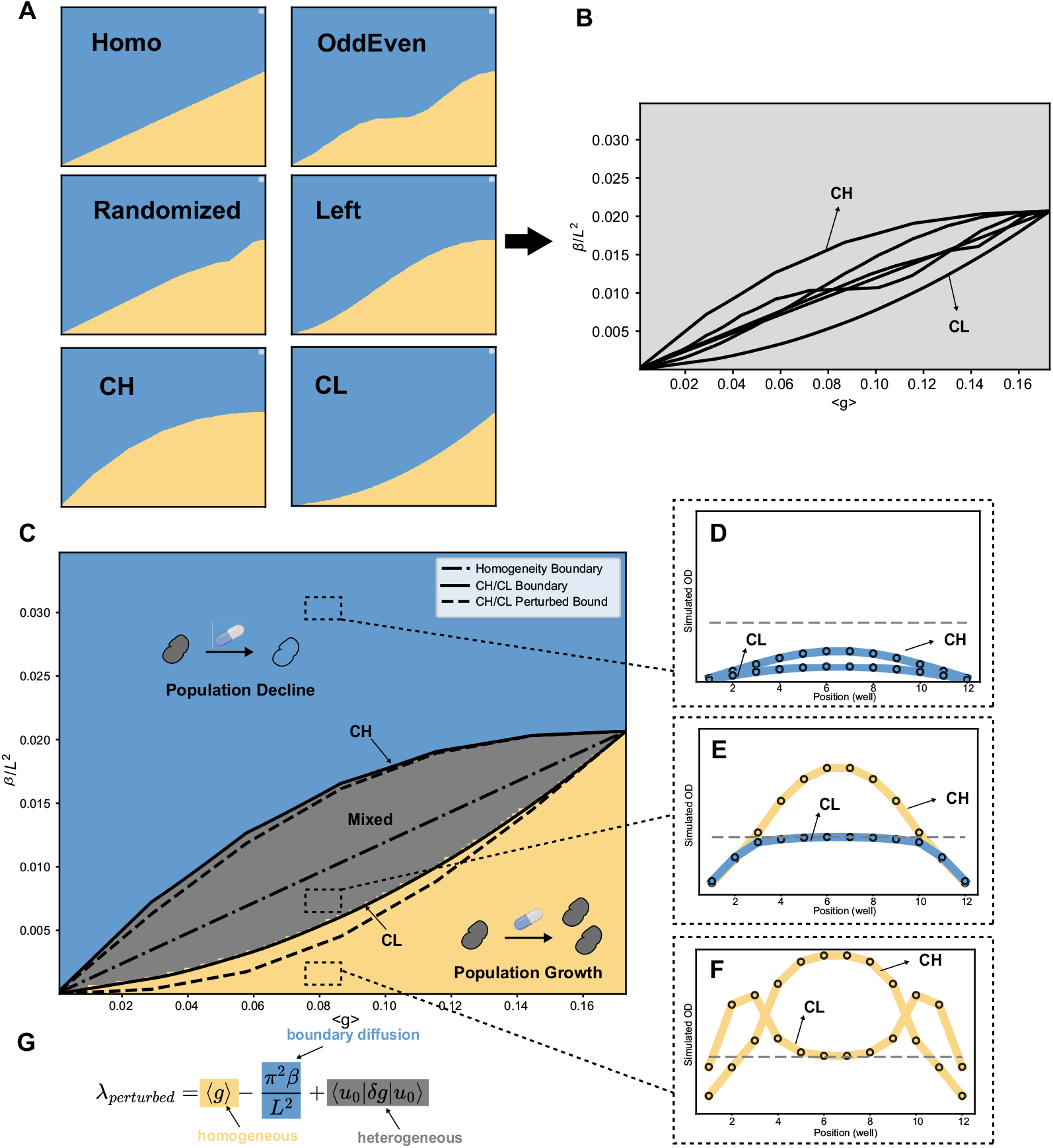
Model validation of the mixed response phase, optimal spatial arrangements CH and CL. **A** and **B**. six different example spatial arrangement strategies imply that CH and CL can possibly be optimal bounds of the emerging mixed phase in the phase diagram of ⟨*g*⟩ and 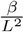. **C**. Phase diagram with new mixed phase by numerically solving the constrained optimization problem eq 4. The solid lines are CH(upper) and CL(lower). They match with the numerical boundary well. The dash-dotted line is homogeneous spatial arrangement. Dotted lines are CH(upper) and CL(lower) by perturbation theory. **D**,**E**,**F**. 3 examples are taken from decline, mixed, growth phase. **G**. The largest eigenvalue by perturbation approximation. It indicates that the diverging responses are incurred roughly by ⟨*u*_0_|*δg*|*u*_0_⟩, an average of spatial growth deviations weighted by square of unperturbed eigenvectors.

To validate the hypothesis that spatial arrangement CH and CL may mitigate population decline most and least effectively, serving as the upper bound and lower bound of the mixed phase, we can translate it into a constrained optimization problem

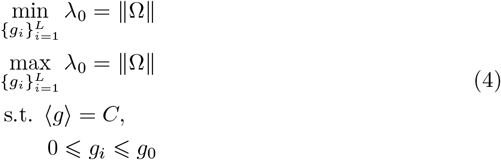

The original equation is discretized with *L* = 12 wells, matching our experimental conditions. The optimization is constrained by a fixed spatially averaged growth rate, while the growth rate at each well *i* is limited to a maximum of the drug-free growth rate *g*_0_ and a minimum of 0, modulated by the drug concentration at each well. The minimizer of the largest eigenvalue equal to 0 corresponds to the lower boundary of the mixed phase, while the upper boundary is determined by finding the maximizer that equals 0 (for more details, see Supplementary Information). When the largest eigenvalue is positive for any spatial arrangement, the population consistently grows. Conversely, when the largest eigenvalue is always negative, the population consistently declines, as shown in Figure 4C. The new mixed phase exhibits different outcomes depending on the spatial arrangements, and it is bounded by CH and CL, as can be proven by the KKT condition. Figure 4D, 4E, and 4F show examples of the decline phase, mixed phase, and growth phase, respectively, where both CH and CL decline, CH grows while CL declines, and both grow. This 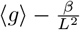 phase diagram not only demonstrates the existence of a mixed-response phase but also indicates that, even under spatial drug heterogeneity, it is possible to ensure bacterial population decline regardless of temporal fluctuations in the spatial drug environment. This can be achieved by driving the population into the decline phase, since any spatial drug arrangement will incur population decline in this parameter region, as shown in Figure 4C.

To better understand this phase diagram and observe that the mixed phase is symmetric around the original homogeneous growth rate boundary line, we apply first-order perturbation theory. The largest eigenvalue can be decomposed into three parts: the homogeneous growth rate ⟨*g*⟩, the boundary diffusion effect 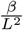, and the heterogeneous effect ⟨*u*_0_|*δg*|*u*_0_⟩ induced by the spatial drug arrangement. Interestingly, the wells can be ranked by the square of their corresponding eigenvector components *u*_0_(*i*)^2^. Given that 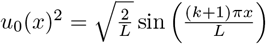 (or 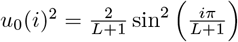 in a discrete version), the wells with the most weights are the center wells. Consequently, when drug-free growth rates are placed at the center, as in CH, *λ*_0_ reaches its maximum, as expected. Thus, the optimal spatial arrangements can be approximately explained by the original eigenvectors driven solely by the boundary diffusion effect. It is also shown that the perturbed eigenvalue is most accurate when the boundary diffusion effect is large (see Supplementary Information).

### Experimental data charts the new mixed phase and empirical boundaries

To validate the theoretical findings that a new mixed phase exists, where different spatial arrangements induce varied dynamic outcomes in addition to the decline and growth phases, we experimentally tested various levels of fraction transfer rate *b* and numbers of drug wells *n*_*D*_, equivalent to tuning migration rate *β* and spatially averaged growth rate ⟨*g*⟩(Figure 5A,B). To avoid the curse of dimensionality from permutations, we focused on the center drug-free wells (CH) and center drug wells (CL) from the new phase diagram, as they define the largest region of the mixed phase and thus lead to the most distinct results (see Figure 4C). As mentioned in the system setup, whether the population declines or not is determined by *λ*_0_. For a fixed number of drug wells (for example, *n*_*D*_ = 6; see Figure 5C), we experimentally increased the fraction of volume transfer *b* to enhance the boundary diffusion effect. We observed that populations in both CH and CL regimes grow when the migration rate/boundary diffusion effect is small. As the boundary diffusion effect increases, the population in the CL regime begins to decline, while the population in the CH regime continues to grow. Eventually, both populations decline when the boundary diffusion effect becomes very large. Our simulation results qualitatively capture these temporal dynamic features (see Figure 5C; also Supplementary Information for a complete simulation illustration). The experimental data(dots) align well with the phase diagram generated by numerically solving the eigenvalues of spatial arrangements CH and CL (see Figure 5B). The mixed phase is still evident in the middle, where CH grows while CL declines.

**Fig 5.**
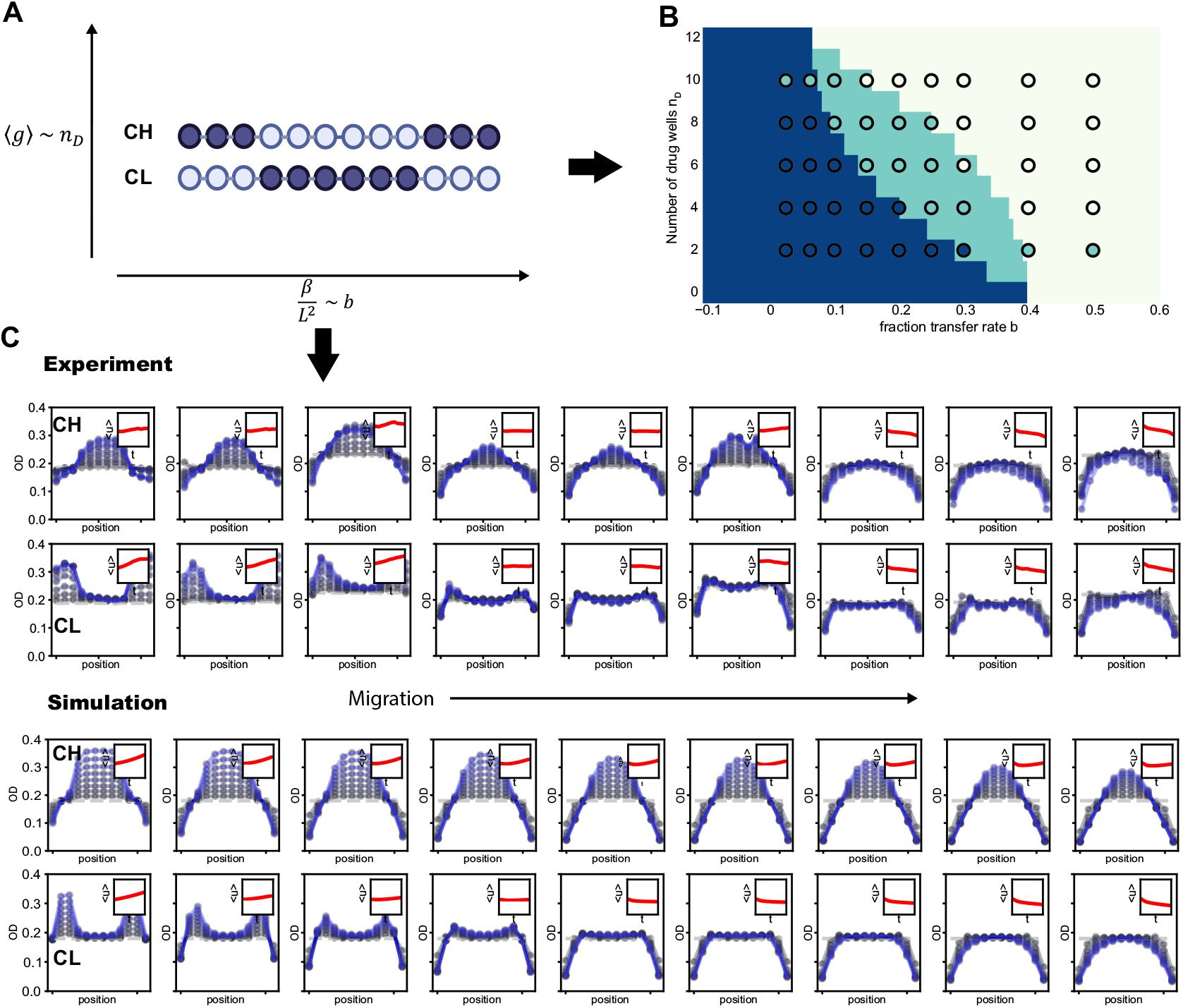
Experimental validation of the new mixed phase. **A**. Increase the number of drug wells *n*_*D*_ to decrease the spatially averaged growth rate ⟨*g*⟩, and increase the fraction of volume transfer *b* to enhance the boundary diffusion effect 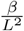. The example shown is *n*_*D*_ = 6(dark color indicates drug wells). **B**. Experimental data (dots) reveals three distinct phases, which qualitatively match the numerical phase diagram. **C**. For the number of drug wells *n*_*D*_ = 6, by increasing the migration rate/fraction of volume transfer *b*, the population responses of both CH and CL shift from growing to declining. The simulation captures the experimental features of these responses and their temporal dynamics. The population under the CH spatial arrangement continues to grow until *b* = 0.3, while the population under the CL arrangement grows only until *b* = 0.1. The inset shows the change in spatially averaged population density, ⟨*u*⟩, over time. The dashed line represents the initial optical density (OD). Grey blue dots and curves correspond to early cycles, while lighter shades of blue indicate later cycles. Each curve is the average of eight technical replicates.

### Spatial effect in other spatially-extended systems

Microbial communities often diffuse and migrate within complex spatial structures. Although the effects of spatial drug heterogeneity, or spatial arrangement on 1D structures with absorbing boundary conditions have been illustrated in previous sections, the interplay between arrangement effects and other spatial structures, beyond the boundary diffusion effect, remains unclear. Here, we hypothesize that spatial drug arrangements can still alter growth dynamics and lead to divergent response outcomes, indicating the existence of a mixed phase, even when other forces drive population decline. To generalize our findings, we designed a ring structure as a periodic condition in our reaction-diffusion model. Although periodic boundary condition has been intensively studied in the context of theoretical ecology, for example for finite one-dimensional or two-dimensional space, or infinite one-dimensional environment [65, 66, 68–70], experimental evidence is rare. The bactericidal drug Ampicillin was applied to specific wells to induce maximum cell lysis [18], creating a deleterious environment or “sink,” while other wells remained drug-free, serving as the bacterial “source.” The interplay between the maximum death rate and the drug-free growth rate, connected by the migration rate *β*, ultimately determines whether the population will grow or decline.

It’s intuitive to expect that at low migration rates, bacteria thrive in drug-free wells with minimal perturbation from drug sink wells. As the migration rate *β* increases, populations with different spatial arrangements diverge into a mixed phase and eventually decline at high migration rates, mirroring what we see with the boundary diffusion effect. Our experimental results with four different spatial arrangements confirm this: as *β* increases, the fraction of population growth conditioned on these arrangements transitions from 1 (growth phase) to 0.25 (mixed phase), and finally to 0 (decline phase) (see Figure 6C). Although different spatial drug arrangements and boundary condition are applied, migration rate still plays a driving factor for bacterial population decline, and the system exhibits similar diverging outcomes.

**Fig 6.**
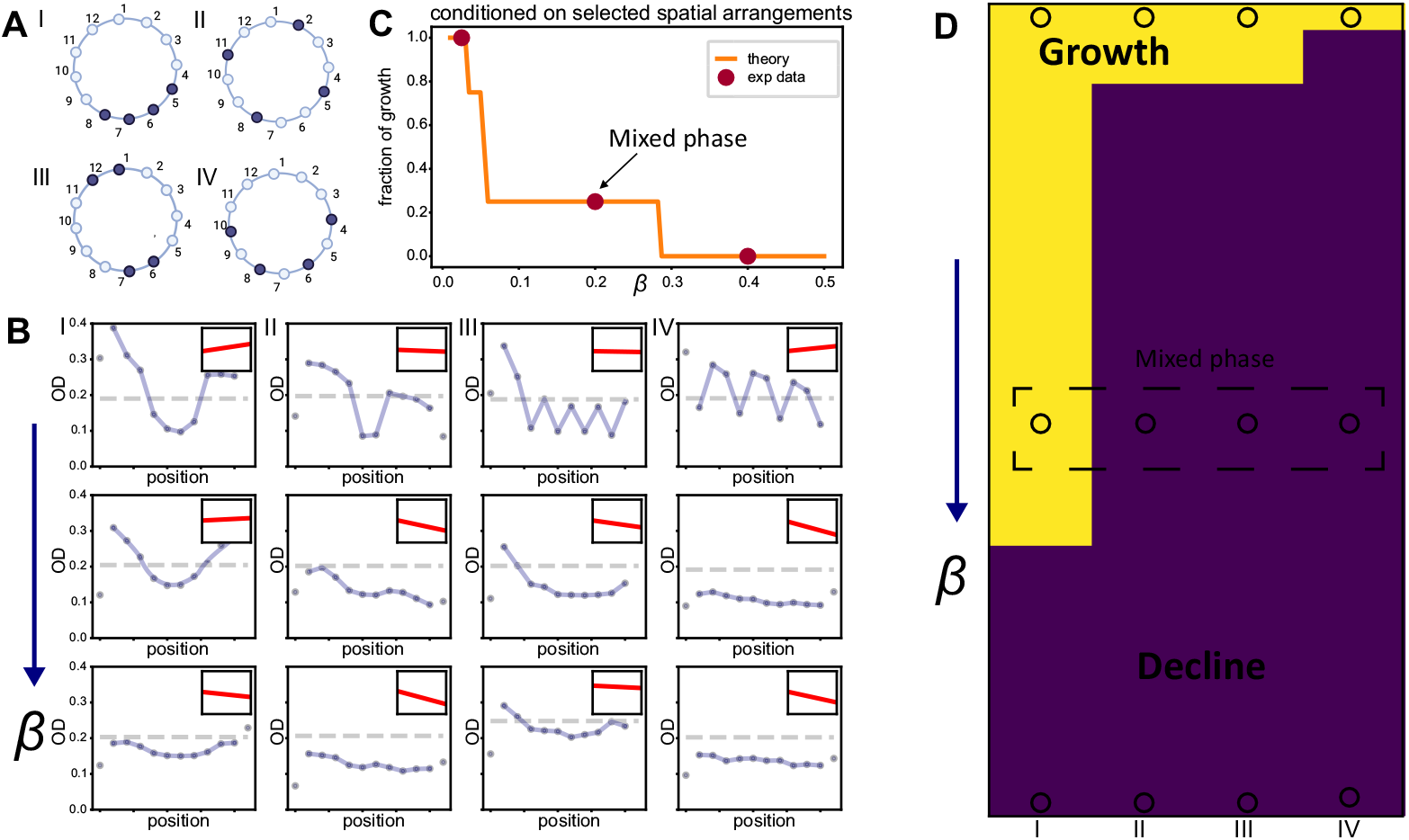
Different spatial drug arrangements on a ring structure induce divergent response outcomes. **A**. Four different spatial arrangements (I, II, III, IV) in a ring structure, where dark color indicates drug wells (*g <* 0), and light color indicates drug-free wells (*g >* 0). **B**. These four spatial arrangements orchestrate different temporal dynamics across three different migration regimes. Dashed line represents the inital OD while blue dots and curves represent endpoint ODs. The inset shows the change in spatially averaged population density, ⟨*u*⟩, over time. **C**. Among the four spatial arrangements, the fraction of growth responses decreases as the migration rate increases; at *β* = 0.2, the fraction is 0.25, indicating the existence of a mixed phase. **D**. The experimental data (dots) aligns with the numerical phase diagram obtained by solving the largest eigenvalue. Yellow indicates growth responses, while dark blue indicates decline responses.

We next examined whether our simplified reaction-diffusion model with a new periodic boundary condition and death rate aligns qualitatively with experimental data. Figure 6B illustrates the temporal dynamics across different migration rates and spatial drug arrangements, closely matching simulation results (see Supplementary Information). Theoretical predictions using the largest eigenvalue criterion also accurately capture population outcomes (see Figure 6D). These agreements validate our model and demonstrate its robustness in more complex scenarios. To explain the emergence of spatial arrangement effects in this new ring structure, the perturbed eigenvalue is calculated. However, due to the equivalence of each well in this spatial structure, the perturbed largest eigenvalue *λ*_*p*_ = ⟨*g*⟩ simply becomes the spatially averaged growth rate, and it fails to give the information of spatial drug heterogeneity. This may necessitate higher-order perturbations or the development of new theoretical tools for further investigation of complex spatial structures.

## Discussion and Conclusion

In this paper, we developed an island-like interconnected experimental system to investigate the effects of spatial drug heterogeneity. We first discovered that simple trade-off relationships—between growth, boundary diffusion effects, and spatial arrangement effects—govern the transition in growth dynamics. Different spatial arrangements of drugs, even with the same spatially-averaged growth rates, lead to divergent bacterial population outcomes, resulting in a mixed response phase. Furthermore, simulation and optimization results identify CH and CL as two optimal spatial arrangements, serving as empirical upper and lower bounds of this mixed phase. This finding is validated through systematic high-throughput experiments. An approximation using perturbation theory explains how spatial drug arrangements alter growth dynamics and lead to different outcomes. It also illustrates that even under spatial drug heterogeneity, it is possible to ensure bacterial population decline regardless of temporal fluctuations in the spatial drug environment, by driving the population from the mixed phase into the decline phase. Further extensions with a ring structure confirm the importance of spatial drug arrangement, showing that spatial drug heterogeneities can incur diverging population responses in other spatially-extended systems.

For the two optimal spatial arrangements with fixed average growth rates, CH and CL, the opposing yet symmetric configurations arise due to the effects of boundary diffusion and the symmetric 1D spatial structure. Interestingly, a similar “positional advantage” has been observed in an evolution experiment conducted in microchannels with the same absorbing boundaries [71], where the dominance advantage of cells at the center position is maximized. This further highlights the impact of boundary conditions. Additionally, spatial structures, or potentially different network configurations, play significant roles in cancer therapies and clinical decisions, often manifesting as star or tree formations [72, 73]. Given that a mixed phase still persists, further investigations are necessary to explore how spatial structures influence the optimal spatial arrangements.

To avoid potential mutations under long-term operations and maintain a consistent environment, the dilution step was omitted in our growth-migration experiments, unlike in the other range expansion experiments doing migration on 96-well plates [57]. While this method is efficient, it may introduce possible drug diffusion, which we minimized its effects in our experiments by choosing appropriate experimental parameters. However, this drug diffusion could be significant in human bodies, where pharmacokinetic-pharmacodynamic (PK-PD) dynamics are at play. In the phase diagram under spatial drug heterogeneity, the ideal theoretical mixed phase region does not align perfectly with the experimental data, which appears narrower. While the model is simplified relative to the complexities of the experimental phenomena and remains powerful enough to qualitatively explain the results, this discrepancy indicates that factors like drug diffusion or other time-varying drug fluctuation, due to natural noise, still exist. Therefore, for those aiming to clinically observe and study the spatial drug heterogeneity effect, it is generally necessary to minimize environmental noise and reduce the drug’s diffusion effect in order to achieve a clearer mixed phase.

Even though, in this study, we focused on the single-species dynamics of wild-type (WT) bacteria, in nature, bacteria often form multi-species communities, and multiple drugs are commonly applied together as part of combination treatments. Our recent work has theoretically explored how antibiotic resistance mutants are selected based on growth dynamics under spatial multi-drug heterogeneities, considering drug interactions [66]. Further experimental and clinical data are needed to validate these findings. Moreover, while our system assumes density-independent exponential growth for pathogens and cancer proliferation [10, 74], different species can have ecological interactions with one another [30, 54, 75–80]. In a community, interactions between species and antibiotics can lead to counterintuitive outcomes [78, 79] and increase the prevalence of antibiotic resistance [75]. Understanding how spatial drug heterogeneity impacts these known behaviors, both at ecological and evolutionary scales, remains an open question that requires further investigation. Recent studies have highlighted the importance of diversity-dependence in dispersal, where interspecific interactions determine spatial dynamics [81]. For instance, recent metapopulation models suggest that quenched disorder in death rates could induce a new phase of global coexistence when considering migration and species interactions [82], offering a promising starting point for further exploration.

Our findings provide insights into clinical migration phenomena, potentially informing pathogen and cancer clearance strategies. Tumors, modeled as complex ecosystems using Generalized Lotka-Volterra (GLV) equations, form heterogeneous metastasis networks influenced by spatial heterogeneity and seeding dynamics [83, 84]. Our work may clarify the role of microbial communities in modulating immune responses and elucidate how spatial heterogeneity and organ-level interventions impact metastasis progression and treatment efficacy [72, 85–88].

The diversity in clinical outcomes necessitates personalized therapies, such as transition therapies for phenotypic switching tumors, and our findings contribute to understanding individual variations [89, 90]. Our spatiotemporal model, capturing spatial drug heterogeneity, can be extended to complex scenarios like metastasis, and integrated with AI for improved mechanistic learning, enhancing predictive accuracy and optimizing treatment strategies [91–96].

To summarize, we have shown that in a deleterious confined environment in which growth rates are unevenly suppressed because of spatial drug heterogeneity, the ecological dynamics and responses are reshaped by the drug spatial arrangements and migration rates. A mixed phase is identified and an optimized center drug strategy can be leveraged to shift response towards decline. The existence of decline phase under spatial drug heterogeneity also indicates a robust pathogen clearance approach regardless of the temporal fluctuation of the spatial drug environment. This highlights the importance of expanding our knowledge of how to tune drug spatial distribution for the potential clinical use, especially in the context of drug treatments and their alarming increased failure of pathogen clearance and cancer metastasis.

## Methods

### Experiment details

*Enterococcus faecalis* strain OG1RF, a Gram-positive bacterium, was cultured overnight in 50% BHI media in 50 ml cell culture tubes. The minimum inhibitory concentration (MIC) of Linezolid was approximately 1.5 µg/ml, and the MIC of Ampicillin was approximately 0.5 µg/ml. Each antibiotic (Linezolid and Ampicillin) was prepared from powder stock and stored at -20 °C. The migration/transfer cycle time was set to 0.25 hours for the homogeneous case and 0.5 hours for the heterogeneous case. Growth rates were determined using a 1:1 ratio of cell culture to a specific drug solution diluted in 50% BHI media. All dilutions and migrations were completed by an OT-2 pipetting robot dispensing into 96-well plates.

### Experimental Protocols

All cultures were grown at 37 °C in 50% BHI media overnight for 18-20h. All experiments were performed in BioLite 96 Well Multidish. For the spatial heterogeneous migration experiment, the same strain was cultivated under two different conditions: 50% BHI media (high growth rate) and 50% BHI media + 8ug/ml Linezolid (low growth rate). Cells were diluted 1:5 with 50% BHI media and grew in a new 15ml cell culture tube for 45 minutes before transferring to the 96-well plates and starting the first migration.(Mix the media with or without drug with cells 1:1 ratio). Cell migrations were carried out along the columns of the plate, in 12-well-long landscapes. Migrations were performed every 15/30 minutes using the Opentron OT-2 robot for ∼ 8 times. Plates were not shaken during growth. Optical densities were measured after every migration cycle in the plate reader. with 600-nm light. To explore more possibilities, we changed the transfer volumes to the neighboring columns during the migration in order to control the migration rate. We transferred 5, 12.5, 20, 30, 40, 50, 60, 80, 100 ul(with the single well transfer rate) to the neighboring columns in different plates. The spatially averaged growth rate is controlled by the number of drug wells. As for boundaries, after discarding a transfer volume, an equivalent volume of media (either containing the drug or drug-free, depending on the drug condition of the boundary wells) was added to maintain a consistent well volume.

### Model details

The one-dimensional Fisher-KPP equation, 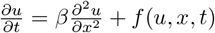, is a well-known equation in ecological and evolutionary dynamics that describes cell growth and range expansion in a spatially varying environment [57, 97]. Here, we consider a special case with linearized growth and fully absorbing boundary conditions (also known as Dirichlet or zero conditions): 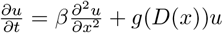, *u*(*x*, 0) = *u_0_, u*(0, *t*) = 0, *u*(*L, t*) = 0, where *L* is the length of the spatial domain. In this scenario, cells can have different growth rates at different positions, but the cell densities at the two boundaries are always zero. If the boundary diffusion effect, *β/L*^2^, is significantly larger than the average growth rate, ⟨*g*(*x*)⟩ (*β/L*^2^ ≫ ⟨*g*(*x*)⟩), the population will decrease. Conversely, if the boundary diffusion effect is much smaller (*β/L*^2^ ≪ ⟨*g*(*x*)⟩), the bacteria population will persist and grow up in the diffusive, deleterious environment. A critical boundary exists where growth and boundary diffusion are balanced when drug concentration is evenly distributed. Under spatial drug heterogeneity, this critical boundary transitions into a critical mixed phase (see SI).

## Acknowledgements

This study was supported by NIH R35GM124875 (KBW), grant GL Proj.2022/0006 awarded by FLAD (TFAF), and the Portuguese Foundation for Science and Technology, FCT (CEECIND/03051/2018, 10.54499/2022.03060.PTDC (EG).

