## Supplementary Information for "Linking spatial drug heterogeneity to microbial growth dynamics in theory and experiment"

#### 1D simplified Fisher-KPP model in a deleterious confined environment

Species invade new territory through a combination of dispersal and local growth. Migration or diffusion in a free environment, like range expansions, are well captured by a simple model incorporating growth and nearest-neighbor dispersal, known as Fisher–Kolmogorov–Petrovsky–Piskunov equation, or Fisher-KPP equation[1, 2],

$$\frac{\partial u}{\partial t} = \beta \frac{\partial^2 u}{\partial x^2} + g(u)u \tag{S1}$$

where $\beta$ is the diffusion or migration rate, $u$ is the cell density, and $g(u)$ is the density-dependent growth rate. To describe the bacteria migration in a confined environment with deleterious boundaries, mathematically, 2 absorbing boundaries are attached and growth rate are linearized for simplicity, by ignoring the intra-species competition

$$\begin{cases} \frac{\partial u}{\partial t} = \beta \frac{\partial^2 u}{\partial x^2} + g(D(x))u \\ u(x,0) = u_0 \\ u(0,t) = 0 \\ u(L,t) = 0 \end{cases} \tag{S2}$$

Now bacteria population are confined within a system of size $L$. $g(D(x))$ is the density-independent growth rate modulated by drug concentration $D$, distributed over $x$. $u_0$ is the initial cell density over space and we here assume it's uniform. To align our 1D experimental system featuring disjointed patches, this modified Fisher-KPP equation can be further discretized, thereby both mimicking metapopulation dynamics typical of many ecological and experimental systems, while also the continuous dynamics frequently assumed in mathematical ecology[1].

##### Transformation to an eigenvalue problem

The Fourier method (also known as separation of variables) is applied below to solve the reaction-diffusion equation. Readers can also try the other methods for their own interest. Consider the following form separating the solution into a product of spatial and temporal components

$$u(x,t) = u_x(x)u_t(t) \tag{S3}$$

Thus the PDE can be re-written as

$$u_x\dot{u}_t = \beta u_x'' u_t + g(D(x))u_x u_t \tag{S4}$$

where $u_x''$ refers to second-order spatial derivative of $u_x$. Divide both sides by $u_x u_t$ and let it equal to a constant $\lambda$,

$$\frac{\dot{u}_t}{u_t} = \beta\frac{u_x''}{u_x} + g(D(x)) = \lambda \tag{S5}$$

Rewrite it as a pair of ODEs with respect to $u_t$ and $u_x$,

$$\frac{\partial u_t}{\partial t} = \lambda u_t, \tag{S6}$$

$$\beta\frac{\partial^2 u_x}{\partial x^2} + g(D(x))u_x = \lambda u_x \tag{S7}$$

where $x \in [0, L]$, $u_t(0) = u_0$, $u_x(0) = 0$ and $u_x(L) = 0$. The solution of the former is $u_t = u_0 e^{\lambda t}$ while the latter defines the eigenvalue problem,

$$\Omega u_k = \left(g(D(x)) + \beta\frac{\partial^2}{\partial x^2}\right)u_k = \lambda u_k. \tag{S8}$$

where $\Omega$ is the operator incorporating growth and migration, $\lambda_k$ is the corresponding kth eigenvalue and $k = 0, 1, 2, \cdots, \infty$. combining the temporal and spatial solutions, the general solution $u(x,t)$ now has the form

$$u(x,t) = \sum_{k=0}^{\infty}\langle u_0|\psi_k\rangle e^{\lambda_k t} \tag{S9}$$

$\langle u_0|\psi_k\rangle = \int_0^L u_0(x)\psi_k(x)dx$ is the coefficient where the initial cell density projected to the eigenbasis. In the long time limit, the cell density are dominated by its largest eigenvalue $u(x,t) \approx \langle u_0|\psi_0\rangle e^{\lambda_0 t}$. When $\lambda_0 < 0$, the whole population goes down and eventually extinct. Thus $\lambda_0$ here can be a good metric to check whether population declines or not. For comparison with the experimental results, we can numerically solve the eigenvalue equation to get $\lambda_0$.

To rescale the spatial coordinate $x$ by L, let $\tilde{x} = x/L$ we have

$$\Omega u_{\tilde{x}} = \left(g(D(\tilde{x}L)) + \frac{\beta}{L^2}\frac{\partial^2}{\partial\tilde{x}^2}\right)u_{\tilde{x}} = \lambda u_{\tilde{x}}. \tag{S10}$$

$\tilde{x} \in [0, 1]$ and now we get our rescaled migration $\frac{\beta}{L^2}$. It incorporates the effect of migration rate $\beta$ and system size $L$. Since in our experiments we either fix migration rate or system size, we won't distinguish rescaled migration or the original migration. For simplicity, let's write back the notation $x$

$$\Omega u_x = \left(g(D(x)) + \frac{\beta}{L^2}\frac{\partial^2}{\partial x^2}\right)u_x = \lambda u_x. \tag{S11}$$

#### Exact eigenvalue solution under spatial drug homogeneity

Assume that the drug concentration is homogeneous over space so $g(D(x)) = g(D) = \langle g\rangle$, the last term is spatially averaged growth rate; in this paper we use this notation to refer growth rate level under homogeneity. Then it becomes a well studied linear Sturm-Liouville system and has an exactly solvable solution because of its integrability. So

$$\lambda_k - \langle g\rangle = -\frac{\beta\pi^2(k+1)^2}{L^2} \tag{S12}$$

And the corresponding eigenvector is $\psi_k = \sqrt{\frac{2}{L}}\sin\left(\frac{(k+1)\pi x}{L}\right)$.This shares the same form as the eigenvectors of pure diffusion equation $\beta\frac{\partial^2 u_x}{\partial x^2} = \lambda u_x$. The population decline criterion is given by the largest eigenvalue $\lambda_0 = \langle g\rangle - \frac{\pi^2\beta}{L^2} < 0$. As mentioned in the main text, the first term is our homogeneous growth rate and the second term is the boundary diffusion effect. They together account for the trade-off.

#### First-order perturbation approximation under spatial drug heterogeneity

Although for a general form of the growth rate $g(D(x))$ under spatial drug heterogeneity, the eigenvalue is not explicitly solvable, we aim to design an approximation that separates the forces driving population growth and decline—namely, the growth rate and boundary effects, as in the homogeneous case. A simple mean field approximation (zero-order approximation) is inadequate here, as it merely yields the spatial average $g_{MF} = \langle g\rangle$, thus neglecting information about spatial arrangement. Instead, we consider a first-order spatial effect using perturbation theory. In our recent work [3], this method demonstrated excellent agreement with numerical eigenvalues, even under multiple-drug resistance evolution with drug interactions and collateral effects. By rewriting the original eigenvalue equation, we can effectively separate the heterogeneous effects.

$$\beta\frac{\partial^2 u_\lambda}{\partial x^2} + \langle g\rangle u_\lambda + \delta g u_\lambda = \lambda u_\lambda \tag{S13}$$

$\delta g = g(D(x)) - \langle g\rangle$ describes the heterogeneous growth deviation and $\langle\delta g\rangle = 0$, modulated by spatial drug heterogeneity. Choose growth with homogeneous drug concentration as the unperturbed system, so we have $u_k = \psi_k = \sqrt{\frac{2}{L}}\sin\left(\frac{(k+1)\pi x}{L}\right)$ as our eigenbasis. The largest eigenvalue $\lambda_0$ now becomes[3]

$$\lambda_0 \approx \lambda_{perturbed} = \langle g\rangle - \frac{\pi^2\beta}{L^2} + \langle u_0|\delta g|u_0\rangle \tag{S14}$$

$u_0$ is the unperturbed eigenvector corresponding to the largest eigenvalue, from the original unperturbed system. This holds when the homogeneous growth rate $\langle g\rangle$, or migration rate $\beta$, is much larger than the heterogeneous growth deviation $\delta g$. So under the high migration or small spatial fluctuation regime, as shown in Figure 5A in the main text, our approximation has small relative errors. It's general for any spatial drug arrangements. We can also elaborate it in another way. Observe that the exact solution $\lambda_0 = \langle u_p|\Omega|u_p\rangle = \langle g\rangle + \langle u_p|\beta\frac{\partial^2 u_\lambda}{\partial x^2}|u_p\rangle + \langle u_p|\delta g|u_p\rangle$, since $\langle u_p|u_p\rangle = 1$. When the perturbed eigenvector $u_p$ close to the unperturbed eigenvector $u_0$, we recover $\lambda_0 \approx \langle g\rangle - \frac{\pi^2\beta}{L^2} + \langle u_0|\delta g|u_0\rangle = \lambda_{perturbed}$. It's illustrated by Figure 5C, showing that a fraction of $u_p$ are close to the unperturbed $u_0$, of all the possible $u_p$s from the boundaries of mixed phase.

As stated in main text, the optimal spatial drug arrangement, CH or CL, is explainable by perturbation theory. To optimize the largest eigenvalue, we need to optimize $\langle u_0|g(x)|u_0\rangle$. Rewrite $\langle u_0|g(x)|u_0\rangle = \frac{2}{L}\int_0^L g(x)\sin^2\left(\frac{\pi x}{L}\right)dx$. So this is a weighted average of $g(x)$ by $\sin\left(\frac{\pi x}{L}\right)$. Noticing that spatially averaged growth rate is fixed, and $\sin\left(\frac{\pi x}{L}\right)$ has the maximum value at center($x = \frac{L}{2}$) and minimum value at boundaries($x = 0, x = L$), so the maximized $\lambda_{perturbed}$ is acquired when we saturate the growth rate at the center as CH strategy. When we saturate the growth rate at boundaries as CL, we get the minimized $\lambda_{perturbed}$. This also holds in a discrete version of our eigenvectors.

**Boundaries of mixed phase by constrained optimization**

For simplicity, a discrete version of simplified Fisher-KPP model is chosen and let $L = 12$wells. To get the boundaries of mixed phase, we can solve the following constrained optimization problems

$$\begin{aligned} &\min_{\{g_i\}_{i=1}^L} \lambda_0 = \|\Omega\| \\ &\max_{\{g_i\}_{i=1}^L} \lambda_0 = \|\Omega\| \\ &\text{s.t. } \langle g\rangle = C, \\ &\qquad 0 \leqslant g_i \leqslant g_0 \end{aligned} \tag{S15}$$

where $\Omega = \beta\mathcal{L} + G$, $\mathcal{L} = \begin{bmatrix} -2 & 1 & & & \\ 1 & -2 & 1 & & \\ & \ddots & \ddots & \ddots & \\ & & 1 & -2 & 1 \\ & & & 1 & -2 \end{bmatrix}$ is the $L \times L$ matrix describing the migration between wells, or discretized Laplacian operator, $G = \begin{bmatrix} \ddots & & \\ & g_i & \\ & & \ddots \end{bmatrix}$ is the growth matrix with ith diagonal entry representing the growth rate at the ith well. The diagonal elements of $G$ are tuned to get optimized largest eigenvalue. $C$ is a constant value. The population growth, mixed, population decline phase are determined when $\lambda_{0,min} > 0, \lambda_{0,max} > 0$, $\lambda_{0,min} < 0, \lambda_{0,max} > 0$, $\lambda_{0,min} < 0, \lambda_{0,max} < 0$. With $\lambda_{0,min} = 0$ the rescaled migration $\beta/L^2$ is solved to determine the lower boundary of mixed phase. Solving $\beta/L^2$ from $\lambda_{0,max} = 0$ gives us the upper boundary of mixed phase.

To prove that CH, CL are the upper and lower boundaries of mixed phase, KKT condition is applied. Here we use the maximization problem as an example. For incorporating inequality constraints, we define the Lagrangian with KKT multipliers $\mu_i$ and $\nu_i$:

$$\mathcal{L}(\lambda_0, g_1, g_2, \ldots, g_L, \mu, \{\mu_i\}, \{\nu_i\}) = -\lambda_0 + \mu\left(\frac{1}{L}\sum_{i=1}^{L} g_i - C\right) + \sum_{i=1}^{L}\mu_i(g_i - g_0) - \sum_{i=1}^{L}\nu_i g_i \tag{S16}$$

The KKT conditions for this problem are:

1. **Stationarity**:

$$\frac{\partial \mathcal{L}}{\partial g_i} = -\frac{\partial \lambda_0}{\partial g_i} + \frac{\mu}{L} + \mu_i - \nu_i = -u(i)^2 + \frac{\mu}{L} + \mu_i - \nu_i = 0$$

2. **Primal Feasibility**:

$$\frac{1}{L}\sum_{i=1}^{L} g_i = C$$

$$0 \leqslant g_i \leqslant g_0 \quad \text{for all } i$$

3. **Dual Feasibility**:

$$\mu_i \geqslant 0, \quad \nu_i \geqslant 0 \quad \text{for all } i$$

4. **Complementary Slackness**:

$$\mu_i(g_i - g_0) = 0, \quad \nu_i g_i = 0 \quad \text{for all } i$$

The second equality from stationarity is derived by Hellmann–Feynman theorem,

$$\frac{\partial \lambda}{\partial g_i} = u^T \frac{\partial \Omega}{\partial g_i} u = u^T e_i e_i^T u = \left(u^T e_i\right)^2 = u(i)^2 \tag{S17}$$

From stationarity we can see that if none of the inequality constraints are active, $\mu_i = \nu_i = 0$, we have $\frac{\mu}{L} = u(i)^2$, for all positions. Since $u(i)^2$ are not uniform due to the boundary condition and spatial drug heterogeneity, so at least we have 1 active inequality constraint. And this gives

$$\begin{aligned} u(i)(g)^2 - \frac{\mu}{L} - \mu_i = 0 \qquad & \text{with} \quad \mu_i > 0, \quad g_i = g_0 \\ u(i)(g)^2 - \frac{\mu}{L} + \nu_i = 0 \qquad & \text{with} \quad \nu_i > 0, \quad g_i = 0 \\ u(i)(g)^2 - \frac{\mu}{L} = 0 \qquad & \text{with} \quad 0 < g_i < g_0 \end{aligned} \tag{S18}$$

And this indicates

$$u(i)^2 > \frac{\mu}{L}, \quad u(j)^2 < \frac{\mu}{L}, \quad u(k)^2 = \frac{\mu}{L} \tag{S19}$$

$i, j, k$ are indices corresponding to drug-free wells, drug wells with full inhibition, and the critical well where the wells are transiting from drug-free wells to drug wells. So to maximize the eigenvalue, we assign $g_0$ at high $u(i)^2$, 0 at low $u(i)^2$. The critical $k$ is when $u(k)^2 = \mu/L$. Figure S1 shows that for $u(i)^2$ from the upper and lower boundaries, $u(i)^2$ are always highest at center and loweset at edge. Thus CH is the optimal spatial drug arrangement for maximized $\lambda_0$ and CL is optimal for minimized $\lambda_0$.

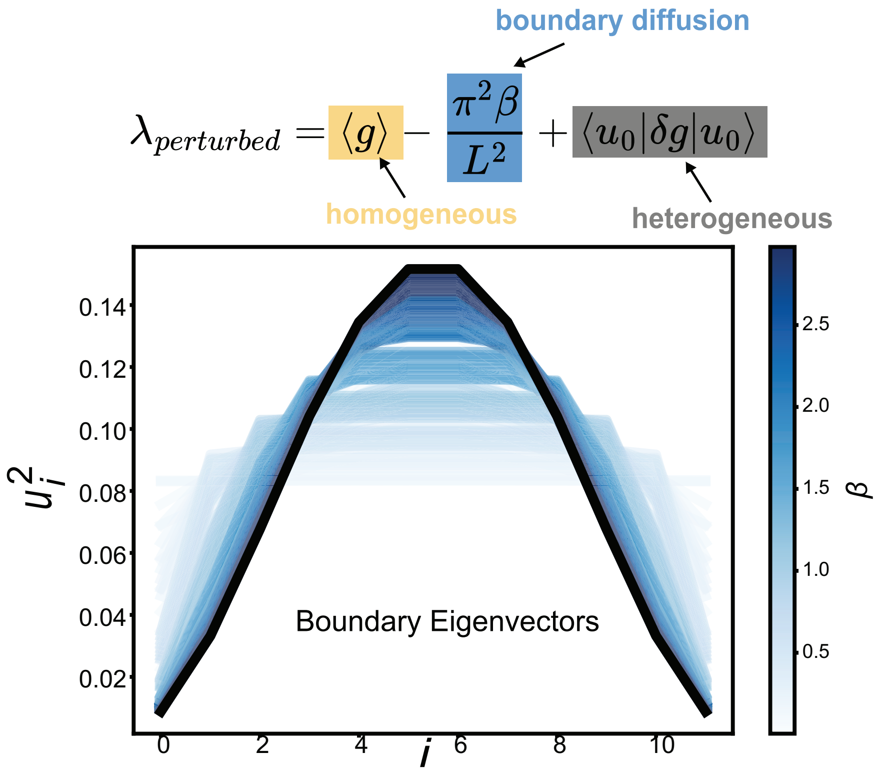


Figure S1: **Unperturbed boundary eigenvectors $u_0(i)$ comparing with eigenvectors $u_0(i)$ under CH/CL spatial arrangements, in a squared form.** As migration rate $\beta$ increases, all eigenvectors from CH and CL spatial arrangements with different spatially averaged growth rates $\langle g \rangle$, are getting closer to $u_0(i)$.

As rescaled migration rate increases, the perturbed eigenvector is closer to $u_0$. From the perturbed eigenvalue $\lambda_{perturbed} = \langle g \rangle - \frac{\pi^2\beta}{L^2} + \langle u_0|\delta g|u_0\rangle$, it's clear that we need to optimize $\langle u_0|\delta g|u_0\rangle = \sum u_0(i)^2(g_i - \langle g\rangle) = \sum \frac{2}{L+1}\sin^2\left(\frac{i\pi}{L+1}\right)(g_i - \langle g\rangle)$. Here we use a discrete version of $u_0$. So, still, because $u_0(i)^2 = \frac{2}{L+1}\sin^2\left(\frac{i\pi}{L+1}\right)$ is maximized at the center and minimized at the edge, we have CH give the maximizer and CL give the minimizer as upper and lower boundaries of the new mixed phase. The KKT condition gives that the optimal $\delta g_i$ values lie at the bounds $-\langle g\rangle$ or $g_0 - \langle g\rangle$, depending on $u_0(i)^2$, and the average constraint ensures the solution is feasible.

Now we have everything to derive the boundaries $\frac{\beta}{L^2}(\langle g\rangle)$. Since $\lambda_0(\mathcal{L}) = -2\frac{\beta}{L^2}(1+\cos\left(\frac{\pi}{L+1}\right))$ in a discrete version, by solving $\lambda_0 = -2\frac{\beta}{L^2}(1+\cos\left(\frac{\pi}{L+1}\right)) + \sum_{i=1}^{L} u(i)^2 g_i = 0$ we get rescaled migration $\frac{\beta}{L^2} = \frac{\sum_{i=1}^{L} u(i)^2 g_i}{2L^2\left(1-\cos\left(\frac{\pi}{L+1}\right)\right)}$, with $\langle g\rangle = C$. This forms the boundary for any spatial arrangement including the homogeneity case, CH, and CL. From the perturbation theory we get

$$\frac{\beta}{L^2} \approx \frac{\sum_{i=1}^{L} u_0(i)^2 g_i}{L^2\left(1-\cos\left(\frac{\pi}{L+1}\right)\right)} = \frac{\sum_{i=1}^{L}\sin^2\left(\frac{i\pi}{L+1}\right) g_i}{L^2\left(1-\cos\left(\frac{\pi}{L+1}\right)\right)} \tag{S20}$$

**Perturbed optimal phase boundaries are always lower**

Rewrite the unperturbed and perturbed eigenvalue function

$$\beta\mathcal{L}\,|u_0\rangle = \lambda_0\,|u_0\rangle\,, \quad (\beta\mathcal{L} + G)\,|u\rangle = \lambda|u\rangle$$

The second equation can be written as $|u\rangle = \frac{1}{\lambda - \beta\mathcal{L}} G|n\rangle$. To eliminate the singularity at $G \to 0$, we define the projection operators

$$P_0 = |u_0\rangle\,\langle u_0| \quad \text{and} \quad Q_0 = 1 - P_0 = \sum_{u\neq q} |q_0\rangle\,\langle q_0|$$

Thus, we have $(1-P_0)\,|u_0\rangle = Q_0\frac{1}{\lambda-\beta\mathcal{L}}G|u\rangle$. The right-hand side can be written as

$$gG|u\rangle \quad \text{where} \quad g = \sum_q \frac{|q\rangle\langle q|}{\lambda - \lambda_0}$$

Therefore, we obtain

$$|u\rangle = |u_0\rangle\,\langle u_0|u\rangle + gG|u\rangle$$

It's known as Lippmann-Schwinger equation.

It is obvious that this equation can be solved iteratively, and its form is

$$|u\rangle = \langle u_0|u\rangle\,(1 + gG + gGgG + \ldots)\,|u_0\rangle$$

$$\langle u|g|u\rangle = \langle u|g|\,\langle u_0|u\rangle\,(1 + gG + gGgG + \ldots)\,|u_0\rangle$$

However, we are interested in understanding eigenvalue perturbation rather than eigenstate perturbation. To solve this, we have

$$\langle u_0 | \beta\mathcal{L} + G | u \rangle = \lambda \langle u_0 | u \rangle$$
$$\text{LHS} = (\lambda_0 + \langle u_0 | G + GgG + GgGgG + \ldots | u_0 \rangle) \cdot \langle u | u_0 \rangle = \text{RHS} = \lambda \cdot \langle u | u_0 \rangle$$

Thus, we have

$$\lambda = \lambda_0 + \langle u_0 | G + GgG + \ldots | u_0 \rangle \quad \text{where} \quad g = \sum_{q \neq u} \frac{|q\rangle\langle q|}{\lambda - \lambda_0}$$

It seems that this is already the general solution, but since $g$ contains the perturbed energy $\lambda$ rather than $\lambda_0$, it still requires a series solution. However, it has been greatly simplified compared to before. Notice that for our specific system, i. $G$ is a diagonal matrix, and ii.$\lambda_{perturbed} - \lambda_0 = \langle u_0 | G | u_0 \rangle > 0$, iii. $u_i^2 > 0$. So apparently $GgG > 0$, and also the $GgGgG, \cdots$ (math induction). so higher order terms are always positive. That's why $\lambda = \lambda_{perturbed} + O(higherorder)$ and $\lambda_{perturbed} < \lambda$.

### Experimental-related design and data analysis

#### Discrete dynamical equation describing the experiment and discretized simplified Fisher-KPP equation

We need to compare the discrete dynamical model that describes the experimental systems with the corresponding discretized form of the 1D Fisher-KPP equation and apply necessary corrections to ensure consistency[1].

According to our experiment protocol, the equation is

$$u_{x,t+\Delta t} = g_{\Delta t}\left(u_{x,t} + b\left(u_{x+\Delta x,t} + u_{x-\Delta x,t} - 2u_{x,t}\right)\right), \tag{S21}$$

where $x$ is the spatial coordinate/well position, $t$ is the cycle number, $b$ is the transfer fraction to a single nearest neighbor, and $g_{\Delta t}(u, x, D)$ describes total growth rate corresponding to the well position $x$ and drug concentration $D$, also the population density $u$, i.e. $g_{\Delta t}(u, x, D)$ is the product of the per capita growth rate by the dose-response curve and the population density. Upon linearization $g_{\Delta t}(u)$ can be written as:

$$g_{\Delta t}(u) = e^{g\Delta t},$$

which corresponds to exponential growth at rate $g$ in the incubation time $\Delta t$.

This is an approximation because, in the experiments, we transfer sequentially from well 1 to well 12 and then in reverse order. However, we approximate this as a simultaneous transfer, thereby ignoring the difference between simultaneous and sequential transfers. Additionally, the dilution effect at the boundary wells may need to be considered. After the transfer, to create the deleterious boundary condition, we still need to take out $bV$ ml liquid with bacteria and add $bV$ ml liquid with fresh media(or with drug). Let's assume that the cell density before this operation is $u$, so the final cell density will be $\frac{1-b}{1+b}u$. Then the effective transfer rate for the boundary wells $b_{edge,eff} = 1 - \frac{1-b}{1+b} \approx 2b$ when $b$ is small.

Using the finite-difference method and discretize the continuous 1D Fisher-KPP equation, we get

$$u_{x,t+\Delta t} = u_{x,t} e^{g_{eff}\Delta t} + \beta_{eff} \frac{\Delta t}{\Delta x^2} \left( u_{x+\Delta x,t} + u_{x-\Delta x,t} - 2u_{x,t} \right) \tag{S22}$$

Comparing eq S21 and eq S22 gives us the effective growth and diffusion rates in the continuous model expressed in terms of experimental parameters as follows:

$$g_{eff} = g,$$

$$\beta_{eff} = b_{eff} \frac{\Delta x^2}{\Delta t} \left(1 + g_{eff}\Delta t\right),$$

where ($\Delta t = 1$ **cycle** $= 0.25$h or $0.5$ h, $\Delta x = 1$ well, and $g = \langle g \rangle$ for the homogeneous case and $g = g(x, D)$ for the heterogeneous case. This results in a 'growth-dependent' effective diffusion rate, which unfortunately leads to an iterative, implicit formulation. For simplicity, we also let $g = g(x, D) \approx \langle g \rangle$ for the heterogeneous case. $b_{eff}$ is set to be $b$ or $2b$ as an approximation, due to the effect of boundary well dilution.

To get the largest eigenvalue, the operator of eigenvalue function can also be discretized as a growth-diffusion matrix and numerically evaluated. Rewrite the eigenvalue function in a discrete version with M positions

$$\beta_{disc} u_{i-1} - 2\beta_{disc} u_i + \beta_{disc} u_{i+1} + g_i u_i = \lambda_i u_i$$

where $i$ is the ith spatial position, $\lambda_i$ is the ith eigenvalue and $u_i$ the ith eigenstate, $k = 1, 2, ...M$; $\beta_{disc} = \beta(M-1)^2/L^2$ In a matrix form we have

$$(\beta_{disc}\mathcal{L} + G)u_\lambda = \lambda u_\lambda$$

where $\mathcal{L} = \begin{bmatrix} -2 & 1 & & & \\ 1 & -2 & 1 & & \\ & \ddots & \ddots & \ddots & \\ & & 1 & -2 & 1 \\ & & & 1 & -2 \end{bmatrix}$ and $G = \begin{bmatrix} \ddots & & \\ & g_i & \\ & & \ddots \end{bmatrix}$. $\mathcal{L}$ is a M by M diffusion matrix and $G$ is a M by M diagonal growth matrix.

#### Response determination

The collective response of the bacterial population can either be growing or declining. In the model the criterion of population growing is the largest eigenvalue $\lambda_0 > 0$. For experiment data, the direct way is to compare the total cell density at the final time $T$ and the total cell density at the beginning,

$$\Delta = \langle OD \rangle_F - \langle OD \rangle_0 = \frac{1}{L}\left(\sum_i^L u_i(T) - \sum_i^L u_i(0)\right)$$

To avoid the boundary well effect when drugs are added to supply both boundary wells, a new threshold $\delta_D < 0$ is introduced, ensuring that $\Delta > \delta_D$. Similarly, when media is added, another threshold $\delta_M > 0$ is applied so that $\Delta > \delta_M$. This criterion was used for generating experimental phase diagrams in the main text.

**Growth estimation**

The model provides three key parameters for tuning: the growth rate $g(D(x))$, the migration rate $\beta$, and the system size $L$ (or 2 effective parameters the growth rate $g(D(x))$ and the boundary diffusion effect $\frac{\beta}{L^2}$). In experimental setups, these correspond to the drug concentration in each well $D(x)$, the transfer rate $b$, and the number of wells $L$, respectively. The transfer rate and number of wells can be predetermined before the experiments commence. Once the drug-modulated growth rates are estimated, the system effectively operates without any free parameters, enabling a direct comparison between the model and experimental results.

For a general estimation of bacterial growth, cultures are incubated under specified conditions for several hours, with the growth rate $g$ calculated as follows:

$$g = \frac{1}{T}\ln\frac{u_f}{u_0} = \frac{1}{T}\ln\frac{OD_{\text{final}}}{OD_0}$$

where $T$ represents the duration of the growth period, $u_f$ is the final cell density, and $u_0$ is the initial cell density, typically measured as optical density (OD). For the growth modulation by the bacteriostatic drug Linezolid (LZD), $T$ is set to 2 or 4 hours. In the case of the bactericidal drug Ampicillin (AMP), bacteria are first grown in a drug-free environment for hours, after which AMP is introduced, leading to growth until a significant decrease in OD is observed. The growth rate $g$ and death rate $d$ are subsequently estimated using the method described in [4]. Results are shown in Figure S2.

**Simulation**

The *PyPDE* Python package is utilized for simulations. *PyPDE* is a Python library designed for solving systems of hyperbolic or parabolic partial differential equations. The simulation closely mirrors the experimental setup. Equation 1 in the main text is discretized into Equation S22 as previously discussed, with a typical system size of $L = 12$. The simulations generally consist of 8 cycles of growth and migration, with parameters selected as outlined earlier.

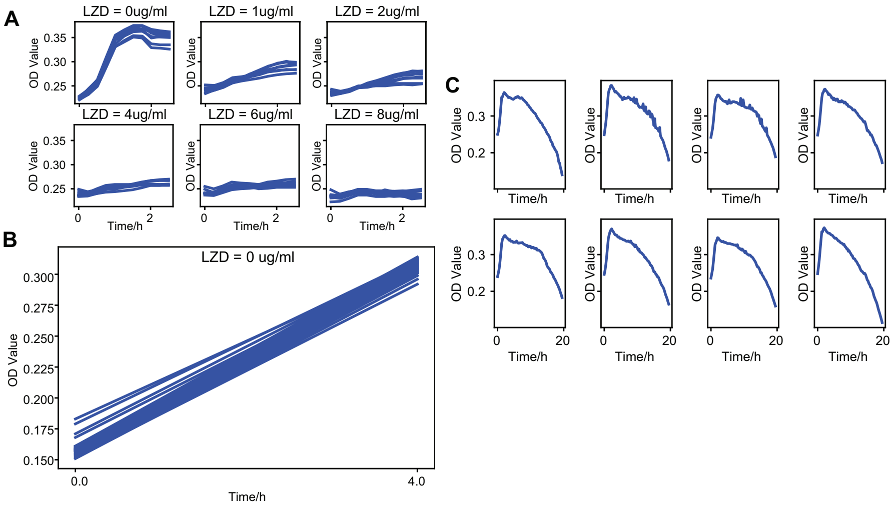


Figure S2: **Growth curves of Homo LZD, Hetero LZD, Hetero AMP. A, B.** Growth curves driven by the bacteriostatic drug LZD, with 8 replicates. In panel A, continuous OD curves under 6 different drug concentrations are plotted. In panel B, only the initial OD and final OD were measured. **C.** Panel C shows 8 individual replicates where bacteria were grown in a drug-free environment, followed by the addition of AMP after hours.

#### Growth validation

Although the growth rate experiments are conducted under the same conditions as the growth-migration experiments, discrepancies may still arise. To further validate our growth rate estimation, we use the CH/CL experimental data as a representative example(See Figure S4, S5) to assess the drug-free growth rate. Reasonable values for $g$ are selected within the range of 0 to 0.32, and the loss is calculated by comparing the data from all 90 experiments with the corresponding simulation results(See Figure S3). Despite the overall high loss, our experimentally determined drug-free growth rate closely aligns with the optimal rate identified from the loss landscape. To illustrate, we selected a random example of $g$ to demonstrate the variation in simulation outcomes(See Figure S6, S7) and compare with the experimental results(See Figure S4, S5). The optimal growth rate determined by the loss landscape was also selected to simulate the dynamics(See Figure S8, S9) and for comparison.

### Simulations and repeated experiments support spatial drug arrangement effect

Detailed evidence has been provided in the main text to demonstrate that different spatial drug arrangements lead to divergent response outcomes. Repeated endpoint experiments and simulations further support this finding. For the

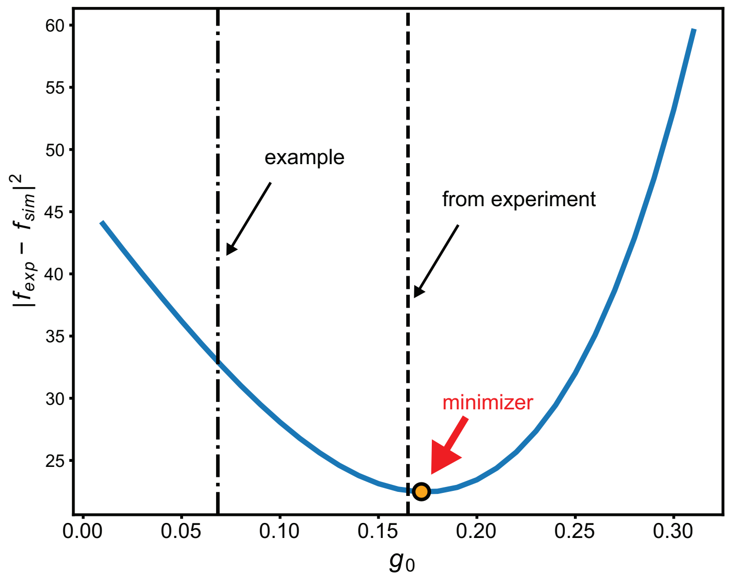


Figure S3: **Loss landscape with drug-free growth rate $g_0$ as the tunable parameter.** The experimentally measured drug-free growth rate is very close to the optimal growth rate shown in this landscape.

initial 6 spatial arrangements, a repeated endpoint experiment was conducted, and despite some discrepancies with Figure 3D in arrangements I and IV, these results, together with the simulation, show strong qualitative agreement (Figure S10A, bottom panel). The simulated temporal dynamics(Figure S10B) also align well with Figure 3B in the main text. In the experimental phase diagram for CH and CL spatial arrangements, endpoints were remeasured with minimal discrepancies(See Figure S11). For different spatial arrangements within a ring structure, the simulation from Figure S12 shows good agreement with Figure 6B.

### Short-term growth rate, long-term growth rate, and largest eigenvalue $\lambda_0$

Although the largest eigenvalue is used in the main text to determine whether the bacterial population is increasing or declining, discrepancies may still arise between the short-term growth rate observed in our short-term diffusion experiments and the largest eigenvalue, which typically represents the long-term growth rate as time approaches infinity. To explore the relationship between the short-term growth rate, long-term growth rate, and largest eigenvalue, we conducted simulations using all CL spatial arrangements to obtain growth rates at different cycle times. The results are presented below(See Figure S13).

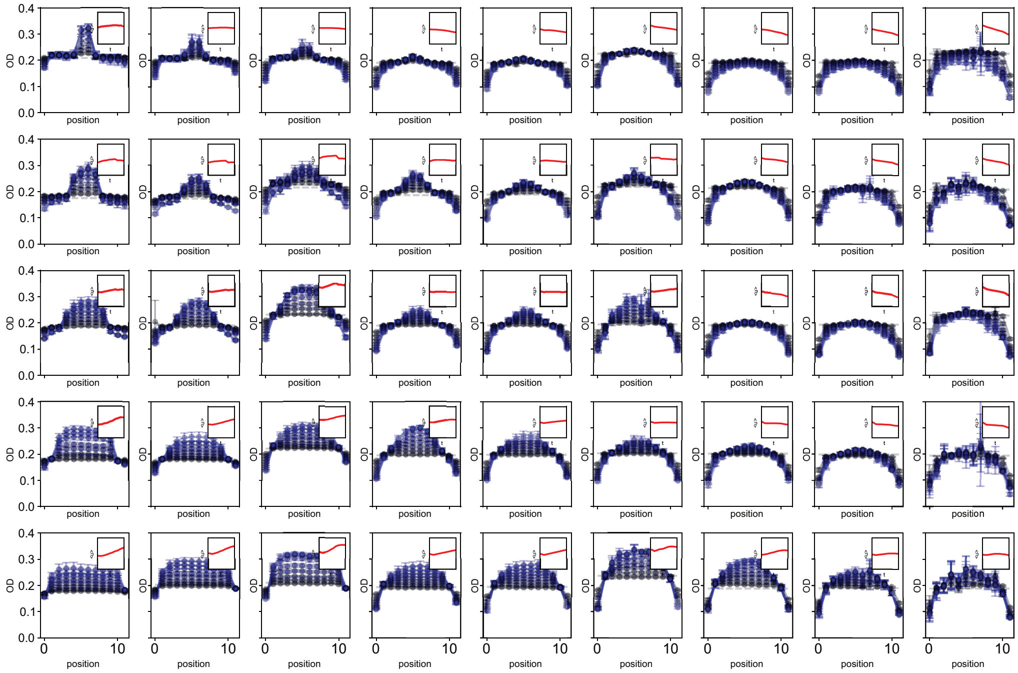


Figure S4: **Experimental result of spatial-temporal dynamics across all conditions under the CH arrangement, with experimentally measured drug-free growth rate $g_{0,exp}$.** The inset shows the change in spatially averaged population density, $\langle u \rangle$, over time. The dashed line represents the initial optical density (OD). Each curve is the average of eight technical replicates, with error bars corresponding to ±1 standard deviation of 8 technical replicates.

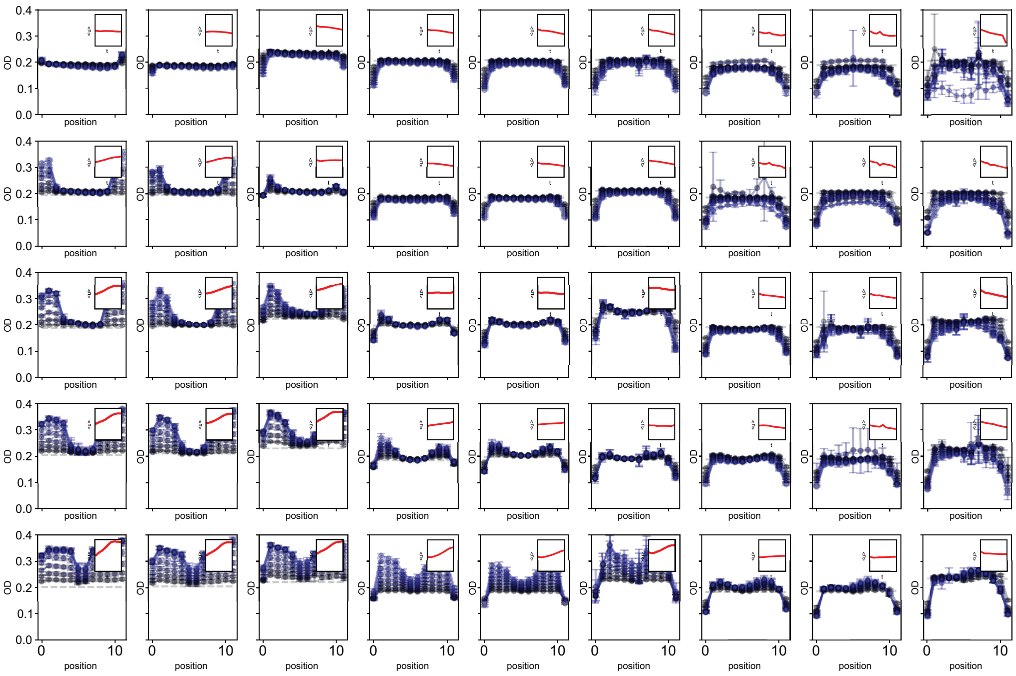


Figure S5: **Experimental result of spatial-temporal dynamics across all conditions under the CL arrangement, with experimentally measured drug-free growth rate $g_{0,exp}$.** The inset shows the change in spatially averaged population density, $\langle u \rangle$, over time. The dashed line represents the initial optical density (OD). Each curve is the average of eight technical replicates, with error bars corresponding to $\pm 1$ standard deviation of 8 technical replicates.

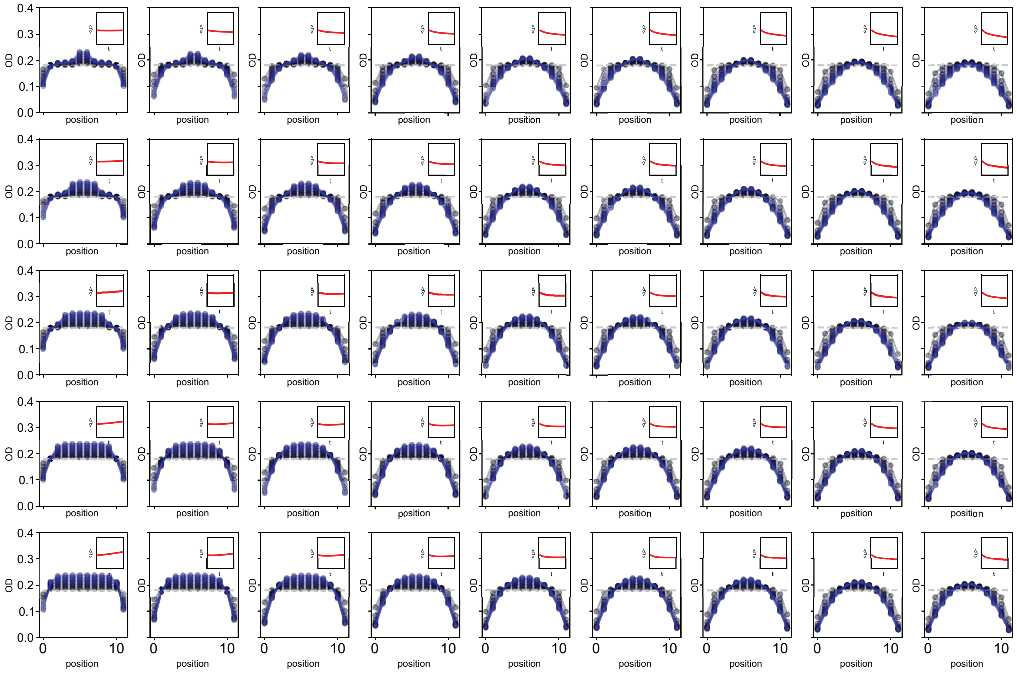


Figure S6: **Simulation of spatial-temporal dynamics across all conditions under the CH arrangement, random sampled drug-free growth rate** $g_{0,rnd}$**.** The inset shows the change in spatially averaged population density, $\langle u \rangle$, over time. The dashed line represents the initial optical density (OD).

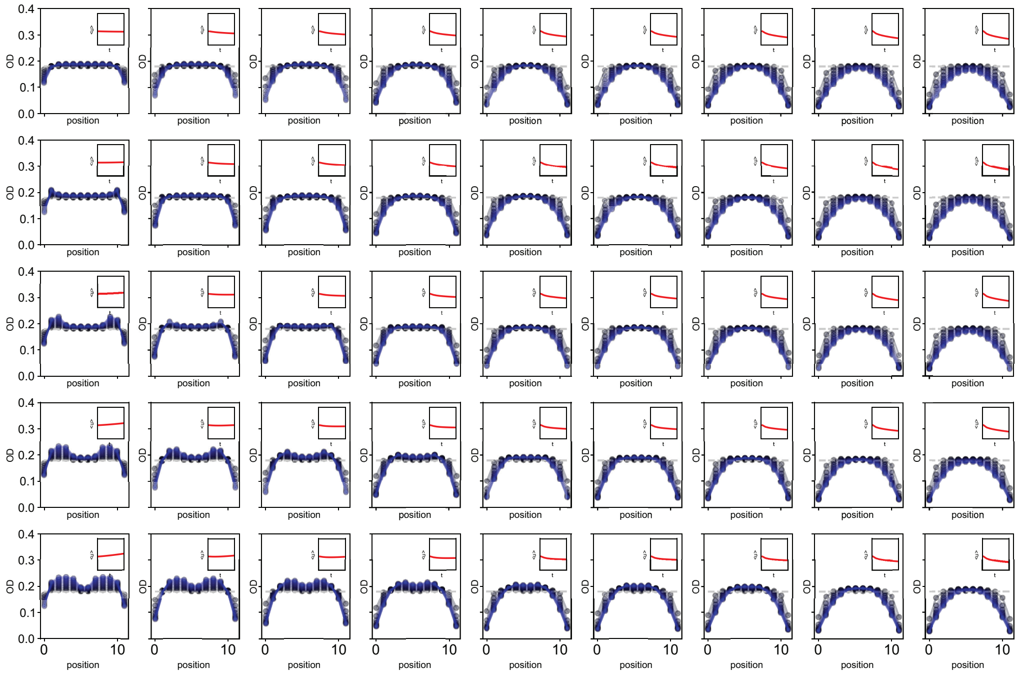


Figure S7: **Simulation of spatial-temporal dynamics across all conditions under the CL arrangement, random sampled drug-free growth rate $g_{0,rnd}$.** The inset shows the change in spatially averaged population density, $\langle u \rangle$, over time. The dashed line represents the initial optical density (OD).

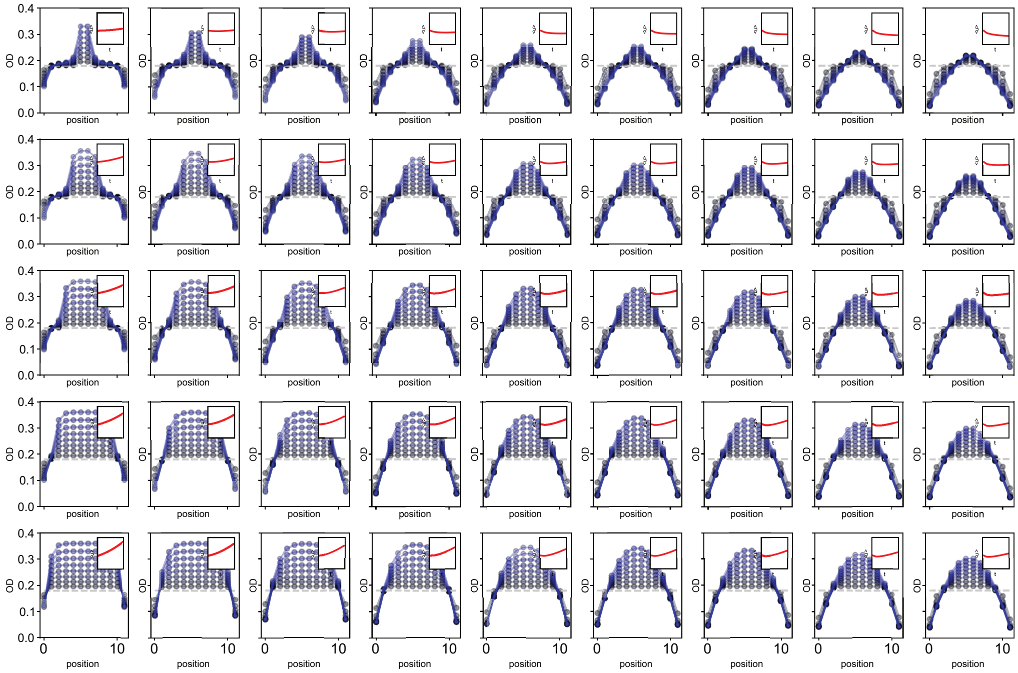


Figure S8: **Simulation of spatial-temporal dynamics across all conditions under the CH arrangement, optimal drug-free growth rate** $g_{0,opt}$**.**The inset shows the change in spatially averaged population density, $\langle u \rangle$, over time. The dashed line represents the initial optical density (OD).

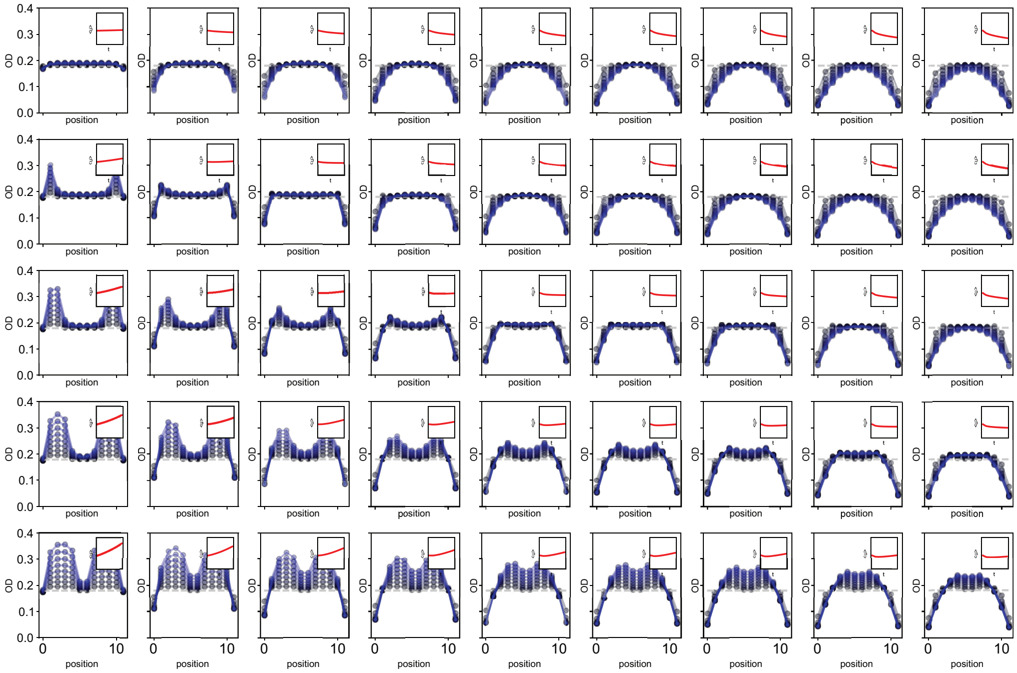


Figure S9: **Simulation of spatial-temporal dynamics across all conditions under the CL arrangement, optimal drug-free growth rate** $g_{0,opt}$ The inset shows the change in spatially averaged population density, $\langle u \rangle$, over time. The dashed line represents the initial optical density (OD).

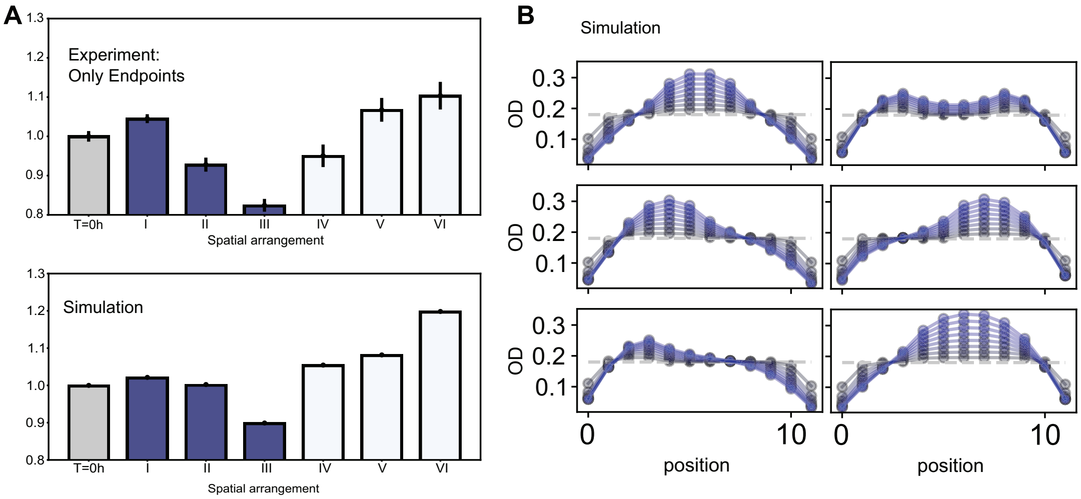


Figure S10: **Repeated endpoint experiment of 6 different spatial arrangements, and the simulation results.** Panel A shows a comparison between endpoint experiment barplot and the result from simulation. Panel B is simulation of temporal dynamics.

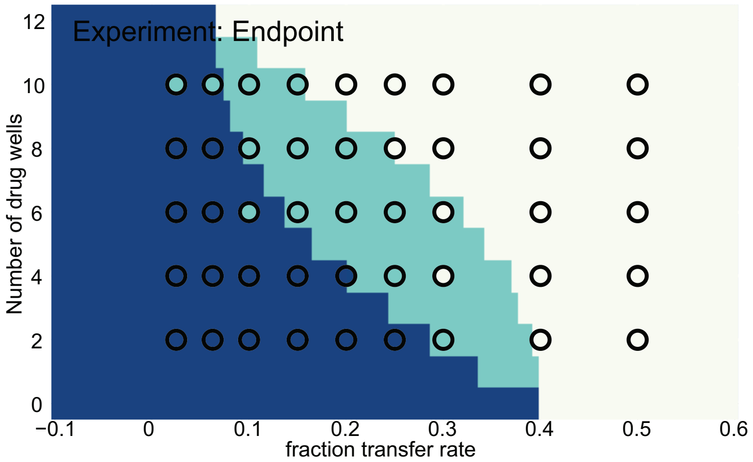


Figure S11: **Experimental phase diagram under CH and CL spatial arrangements, only endpoint.**

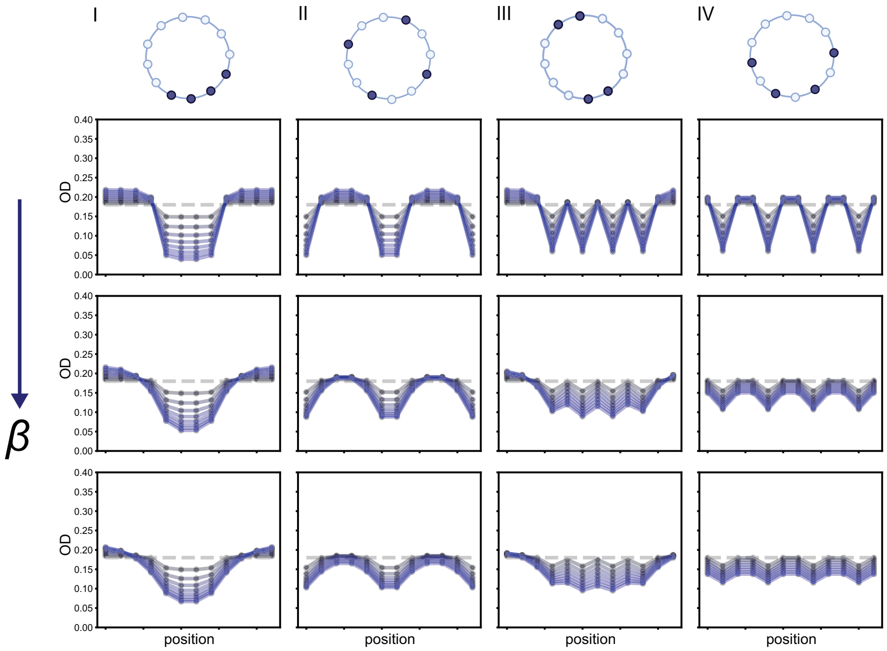


Figure S12: **Simulation of temporal dynamics with a ring structure, for 4 spatial arrangements in 3 different migration regimes.**

Additionally, we compared the perturbation approximation with the numerically solved largest eigenvalue(See Figure S14).

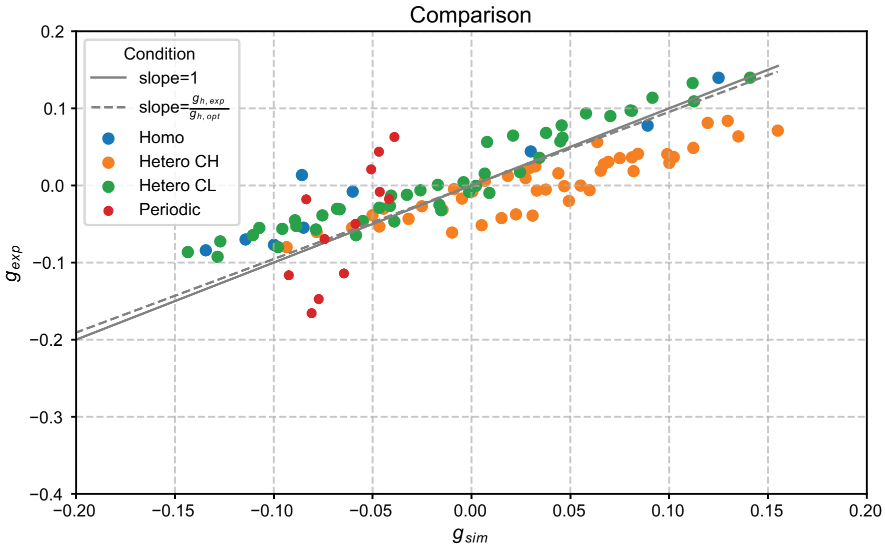


Figure S13: **Comparison between simulated short-term growth rate and experimental short-term growth rate.** Growth rates under different conditions - Homo, Hetero CH, Hetero CL, Periodic - are scattered. The solid line has a slope of 1 while the dashed line is a ratio between growth rates estimated from experiments and from the optimal point of the loss landscape. The experimental short-term growth rates match with simulated short-term growth rates well.

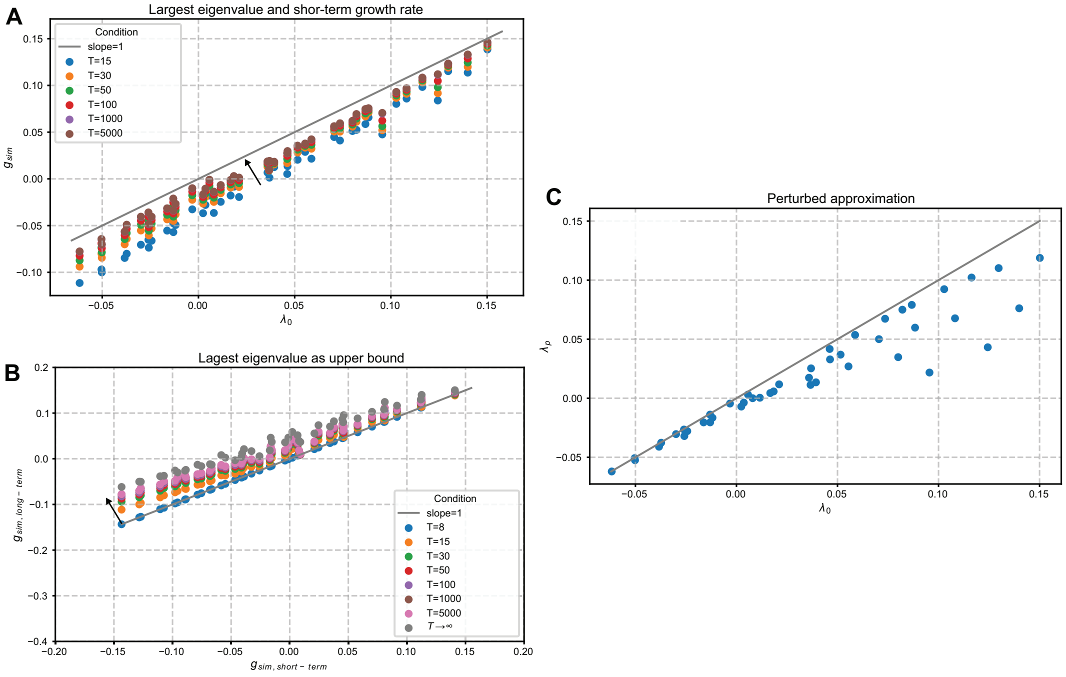


Figure S14: **Comparison between short-term growth rate, long-term growth rate, largest eigenvalue, and perturbation approximation.** Panel A shows that as cycle times increase, the short-term growth rate is asymptotically approaching the largest eigenvalue - the largest eigenvalue is effectively the long-term growth rate after infinite time; Panel B shows that short-term growth rate is always lower than long-term experiments; Panel C shows that the perturbation approximation matches with real eigenvalues well when the original eigenvalue is small.

[4] A. J. Lee, S. Wang, H. R. Meredith, B. Zhuang, Z. Dai, and L. You, “Robust, linear correlations between growth rates and $\beta$-lactam–mediated lysis rates,” *Proceedings of the National Academy of Sciences*, vol. 115, no. 16, pp. 4069–4074, 2018.
